# Phase Separation Potential of Marsupial RSX RNA Reveals Convergent Evolution of X-Chromosome Inactivation Mechanisms

**DOI:** 10.64898/2026.09.18.752704

**Authors:** Ajay Kumar Danga, Camilla Canale, Riccardo Delli Ponti, Gian Gaetano Tartaglia, Andrea Cerase

## Abstract

**Background:** X-chromosome inactivation (XCI) evolved independently in eutherian and marsupial mammals, where it is orchestrated by the unrelated long non-coding RNAs Xist and RSX, respectively. Xist organizes a repressive nuclear compartment through multivalent RNA–protein interactions, but whether RSX exploits similar biophysical principles remains unknown. A recent paper has identified *bona fide* RSX interacting proteins.

**Results:** We integrated proteome-scale RNA–protein interaction prediction, experimental validation, phase-separation propensity analysis, functional annotation and comparative RNA-structure modelling to characterize the RSX interaction landscape. Using *cat*RAPID, we ranked 1,168 RNA-binding proteins from the native *Monodelphis domestica* proteome. Predictions were significantly enriched for experimentally identified RSX interactors, with 4.85-fold enrichment among the top 50 candidates (*P* ≈ 1.6 × 10⁻⁵), increasing to approximately eightfold for proteins shared by the experimental RSX and Xist interactomes (*P* ≈ 2 × 10⁻⁶). Among 30 high-confidence RSX interactors, 13 were experimentally supported, 17 were previously unrecognized candidates and 17 exhibited high phase-separation propensity. The network was enriched in ribonucleoprotein granules and nuclear bodies and converged on m⁶A regulators and SR-family splicing factors. Comparative modelling detected no conserved secondary or tertiary architecture between RSX and Xist.

**Conclusions:** RSX and Xist appear to have converged not through RNA sequence or global structure, but through recruitment of related, condensation-prone protein networks. These findings identify interaction-network and biophysical convergence as a potential principle of lncRNA-mediated chromosome regulation and provide testable candidates for determining whether RSX establishes a condensate-like compartment on the marsupial inactive X.

## Background

X-chromosome inactivation (XCI) is a paradigm of epigenetic gene regulation that ensures dosage compensation between XX females and XY males in mammals [1,2]. In eutherians (placental mammals), XCI is orchestrated by the long non-coding RNA (lncRNA) Xist (X-inactive specific transcript), a 17-19 kb transcript that coats the inactive X chromosome (Xi) in cis and recruits silencing complexes to establish facultative heterochromatin [3,4]. Recent studies have shown that Xist functions through liquid-liquid phase separation (LLPS), forming biomolecular condensates that spatially compartmentalize the Xi and exclude transcriptional machinery [5–7]. Xist achieves this by recruiting LLPS-competent RNA-binding proteins (RBPs) enriched in intrinsically disordered regions (IDRs), which drive formation of membraneless nuclear bodies concentrating repressive chromatin modifiers [8–10].

In contrast, marsupials (metatherians) achieve XCI via a distinct lncRNA, RSX (RNA on the silent X), which shares no sequence homology with Xist [11,12], but share conservation at k-mer composition level [13]. RSX is a 27 kb polyadenylated transcript expressed exclusively from the Xi in female marsupials, representing a striking case of convergent evolution [14]. Despite their independent origins approximately 160-180 million years ago, both RSX and Xist achieve chromosome-wide silencing [15]. This convergence raises a central question: do RSX and Xist utilize similar biophysical mechanisms, particularly LLPS, to accomplish XCI?

Several observations suggest mechanistic parallels between RSX and Xist. Both lncRNAs coat the Xi, recruit chromatin-modifying complexes, and promote heterochromatin formation through histone modifications [16,17]. However, marsupial XCI exhibits unique features, such as tissue-specific incomplete silencing and variable stability across cell types [18]. Critically, whether RSX forms phase-separated nuclear condensates analogous to Xist condensates remains experimentally uncharacterized.

A further parallel between Xist and potential RSX mechanisms involves N⁶-methyladenosine (m⁶A), the most abundant internal modification of mammalian mRNA and lncRNA. m⁶A modification of Xist itself has been shown to promote its association with chromatin-silencing machinery through the nuclear m⁶A reader YTHDC1, directly coupling epitranscriptomic regulation with Xi silencing [19]. The m⁶A writer complex (METTL3–WTAP) and its cognate readers YTHDC1 are LLPS-competent proteins that partition into phase-separated nuclear condensates [20], while the cytoplasmic readers YTHDF1-3 similarly undergo LLPS but in cytoplasmic mRNA-decay and stress-granule-associated compartments [20]. Whether analogous m⁶A-based regulation operates in marsupial XCI through RSX remains unexplored, but represents a compelling mechanistic hypothesis given the conservation of the m⁶A machinery across mammals. If RSX functions through LLPS, it would represent a remarkable example of convergent biophysical mechanisms solving the same regulatory challenge through evolutionarily independent molecular components.

LLPS has emerged as a central organizing principle of nuclear architecture, creating dynamic compartments that concentrate specific factors while excluding others [21,22]. Phase-separated RNA-protein condensates such as nucleoli, nuclear speckles, paraspeckles, and stress granules arise through multivalent interactions between IDR-containing RBPs and scaffold RNAs [23,24]. Xist exemplifies this principle, containing multiple repeat regions that recruit RBPs with high LLPS propensity and low-complexity domains [25,26]. These interactions promote the assembly of Xist condensates that exclude RNA polymerase II and enrich silencing factors including SPEN, SHARP, and Polycomb repressive complexes [10,27].

Advances in computational prediction of RNA-protein interactions and LLPS propensity have enabled systematic exploration of RNA-mediated condensate formation [28,29]. *cat*RAPID predicts RNA-protein binding based on physicochemical features such as secondary structure, hydrogen bonding, and van der Waals interactions [28,30]. catGRANULE 2.0 ROBOT extends this framework with significantly improved accuracy, estimating phase separation propensity from protein sequence characteristics including IDRs, low-complexity domains, and prion-like regions [31,32]. Integrating these complementary approaches allows comprehensive identification of LLPS-competent RBPs potentially driving RNA condensate formation.

Here, we apply an integrated computational framework combining catRAPID omics v2.0 [28] and catGRANULE 2.0 [31] to characterize the RSX interactome against the native *Monodelphis domestica* RBP proteome. Critically, we validate our predictions statistically against an independent experimental RSX interactome [33], demonstrating significant enrichment at multiple thresholds. We identify 30 high-confidence interactors at catRAPID ranking score ≥ 0.50, of which 17 exhibit high LLPS propensity by catGRANULE 2.0 scoring. Together, these findings support a model in which RSX drives chromosome-wide silencing through condensate formation, suggesting that LLPS represents a conserved biophysical strategy for dosage compensation across mammalian evolution.

## Results

### Predictions of the experimental RSX interactome

*cat*RAPID omics v2.0 predictions of RSX-protein interactions were generated against the native *Monodelphis domestica* RBP proteome (1,168 proteins, submitted as 9 proteome chunks; see Methods). To assess the predictive accuracy of our *cat*RAPID analysis, we validated the predictions against an independently published experimental RSX interactome [33]. The prediction universe comprised 1,140 proteins from the merged *ca*tRAPID output files, after deduplication of unique gene symbols. Of these, 47 proteins matched experimentally identified RSX interactors, establishing a baseline probability of 4.1% (47/1140) for recovering an experimental hit at random. This low baseline reflects the fact that experimentally validated interactors represent a small, stringent subset of the much larger pool of computationally predicted RNA-binding proteins.

Proteins were ranked by mean catRAPID ranking aggregated at the gene level. Among the top 50 predicted proteins, 10 were experimentally validated (expected by chance: 2.06; see Methods), representing a 4.85-fold enrichment (hypergeometric p ≈ 1.6 × 10⁻⁵). Among the top 100 predictions, 13 were experimentally validated (expected: 4.12), a 3.15-fold enrichment (p ≈ 9.5 × 10⁻⁵). For proteins detected in both RSX and Xist experiments, enrichment in the top 50 predictions reached ∼8-fold (observed: 8, expected: ∼1; p ≈ 2 × 10⁻⁶) **(Figure 1)**.

**Figure 1.**
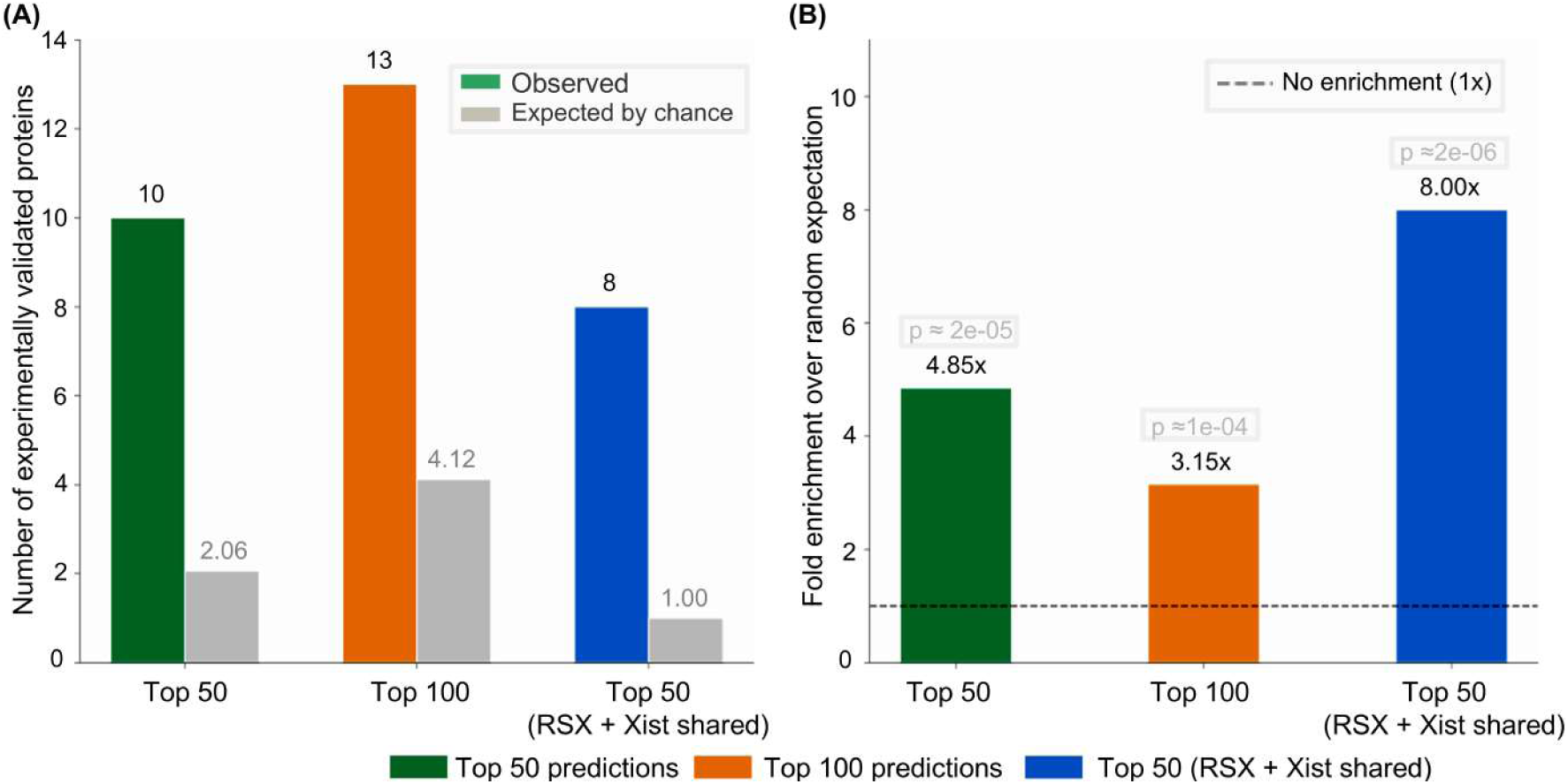
Statistical validation of catRAPID omics v2.0 predictions against the experimental RSX interactome. **(A)** Bar chart showing observed versus expected numbers of experimentally validated RSX-interacting proteins among the top 50 predictions, top 100 predictions, and top 50 predictions restricted to proteins shared between RSX and Xist experimental datasets. Expected values (grey bars) were calculated based on the baseline probability of 47/1,140 ≈ 4.1% derived from the full prediction universe. Observed values (coloured bars) represent proteins experimentally identified in McIntyre et al. (Genome Biology, 2024; DOI: 10.1186/s13059-024-03280-0). **(B)** Fold enrichment of experimentally validated proteins over random expectation for the same three prediction subsets. Fold enrichment values are 4.85× (top 50; hypergeometric p ≈ 1.6×10⁻⁵), 3.15× (top 100; p ≈ 9.5×10⁻⁵), and ∼8× (top 50, RSX+Xist shared; p ≈ 2×10⁻⁶). The dashed horizontal line indicates no enrichment (1×). Statistical significance was assessed using the hypergeometric test. The threshold of catRAPID ranking score ≥ 0.50 defines the 30 high-confidence interactors used in subsequent analyses. Expected values represent the number of experimental hits predicted by chance alone (baseline rate of 47/1,140 experimentally validated proteins × N), against which observed overlap was tested by hypergeometric enrichment.

The strongest enrichment occurred at a threshold of catRAPID ranking score ≥ 0.50, defining the 30 high-confidence interactors used in subsequent analyses **(Figure 1)**. Functionally, these high-confidence interactors are enriched for RNA regulatory processes, including RNA splicing (SRSF1, SRSF2, SRSF3, SRSF5, SRSF7, SRSF9, TRA2A, U2AF1; 26 %), RNA processing and RNA binding (ELAVL1, FUS, FMR1, AGO1; 13 %), and the m⁶A RNA modification machinery (METTL3, WTAP, YTHDC1, YTHDF1-3; 19 %). Notably, SRSF3, SRSF5, and SRSF9 are also reported as Xist interactors in independent studies (**Table 2**), and Trotman, Calabrese and colleagues recently showed that Xist Repeat A assembles SR proteins to recruit SPEN and drive silencing [34]. Our identification of SRSF2, SRSF3, SRSF5, SRSF6, and SRSF9 as high-confidence RSX interactors suggests an analogous SR-protein-mediated assembly on RSX.

Having established that top-ranked predictions are significantly enriched for experimentally validated interactors, we next characterized the full predicted RSX interactome to identify additional candidate partners and examine its broader functional and biophysical properties.

### catRAPID omics v2.0 analysis of the RSX interactome against the *Monodelphis* ***domestica*** proteome

To ensure biological consistency, we performed catRAPID omics v2.0 predictions of RSX RNA-protein interactions using the native *Monodelphis domestica* RBP proteome (1,168 unique proteins), submitted as 9 validated chunks to ensure complete proteome coverage **(Figure 2; Supplementary Figure 1; Supplementary Table 1)**. This approach supersedes prior analyses based on the mouse proteome — such as those predicting Xist–RBP interactions using human mouse screens [35] — and directly addresses the biological question of which marsupial proteins interact with RSX in its native cellular context.

**Figure 2.**
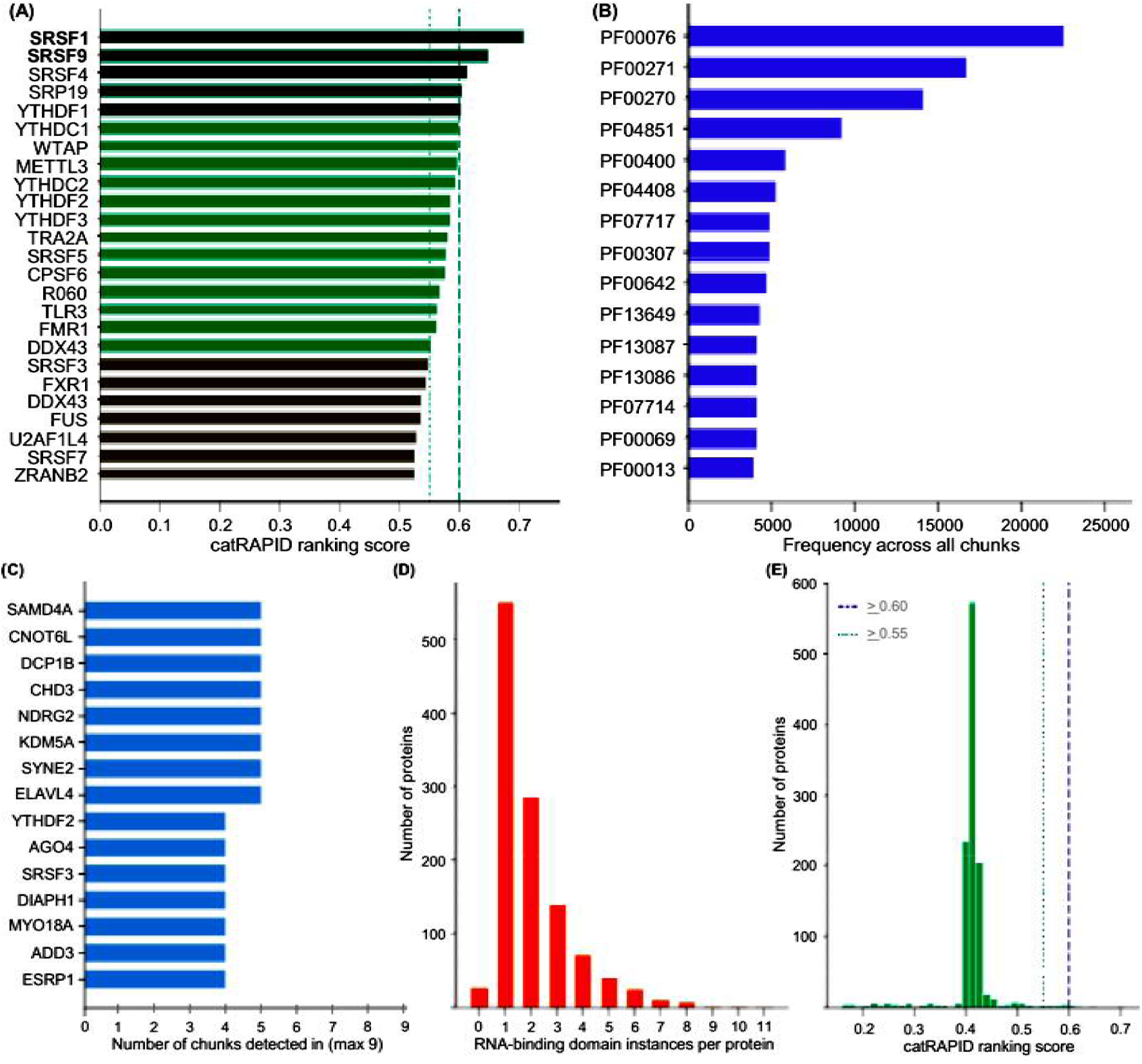
Summary of catRAPID omics v2.0 interaction predictions for RSX lncRNA against the *Monodelphis domestica* proteome across 9 proteome chunks. **(A)** The 25 top-ranked predicted RNA-binding proteins ordered by mean *cat*RAPID ranking score aggregated across all 9 proteome chunks. SRSF1 (0.707) and SRSF9 (0.648) emerged as the highest-confidence interactors. Bars are colour-coded by confidence tier: high (green, ≥ 0.60), moderate (teal, ≥ 0.55), and lower (grey, < 0.55). Dashed and dotted vertical lines indicate the 0.60 and 0.55 score thresholds, respectively. **(B)** The 15 most frequently occurring Pfam RNA-binding domain families across all predicted interactors aggregated over 9 chunks. PF00076 (RNA Recognition Motif, RRM; n = 22,540), PF00271 (Helicase C-terminal; n = 16,660), and PF00270 (DEAD-box helicase; n = 14,112) were the three most prevalent domain families. **(C)** The 15 proteins most consistently detected across proteome chunks, ranked by number of chunks in which they were identified (maximum possible = 9). SAMD4A, CNOT6L, DCP1B, CHD3, NDRG2, KDM5A, SYNE2, and ELAVL4 were each detected in 5 of 9 chunks. **(D)** Distribution of RNA-binding domain instances per predicted interactor protein. The majority of proteins carried 1–2 domain instances (n = 552 and n = 288, respectively), with a minority harbouring ≥ 5 instances. **(E)** Frequency distribution of mean catRAPID ranking scores across all predicted interactor proteins. The distribution is right-skewed, with most proteins scoring between 0.3–0.55, and a distinct high-confidence tail above 0.60.

Aggregation of predictions identified SRSF1 (*ca*tRAPID ranking score = 0.707; see Methods) and SRSF9 (0.648) as the highest-confidence interactors, followed by SRP54, SRP19, YTHDF1, YTHDC1, WTAP, METTL3, YTHDC2, YTHDF2, TRA2A, SRSF5, CPSF6, and RO60 all scoring ≥ 0.55 **(Figure 2A)**. The predominance of m⁶A reader/writer proteins (YTHDC1, YTHDF1/2, METTL3, WTAP) and splicing regulators (SRSF1, SRSF5, SRSF9, TRA2A) among high-confidence interactors points to strong enrichment for m⁶A regulatory and splicing machinery.

Analysis of Pfam domain composition – a systematic classification of conserved protein domains and structural/functional families curated in the Pfam database – was performed across all predicted interactors to characterize the structural basis of their RNA-binding capacity. Each domain family in Pfam corresponds to a recognizable sequence/ structural signature associated with a specific molecular function (e.g., direct RNA contact, ATP-dependent unwinding), so tallying domain occurrence across the interactome indicates which RNA-binding modalities are most represented among predicted RSX partners. This analysis revealed that the RNA Recognition Motif (PF00076; n = 22,540 occurrences), Helicase C-terminal domain (PF00271; n = 16,660), and DEAD-box helicase domain (PF00270; n = 14,112) were the three most prevalent RNA-binding domain families **(Figure 2B)**. The RPM is the most common single-stranded RNA-binding fold in eukaryotes and is frequently found in multiple copies within splicing factors and hnRNPs, while the DEAD-box/Helicase C-terminal domains together define ATP-dependent RNA helicases that remodel RNA structure and RNP composition. Their dominance is consistent with RSX engaging a broad spectrum of canonical RNA-binding proteins rather than a narrow, specialized subset. Most predicted interactor proteins carried 1–2 RNA-binding domain instances, though a subset harboured up to 11 domain instances per protein **(Figure 2D)**; proteins with multiple domain copies are generally capable of multivalent RNA engagement, a property mechanistically linked to RNA-driven condensate assembly.

Cross-chunk detection frequency analysis identified ELAVL4, SYNE2, KDM5A, NDRG2, CHD3, DCP1B, CNOT6L, and SAMD4A as the most consistently predicted interactors, each detected across 5 of 9 proteome chunks **(Figure 2C)**, indicating robust and reproducible enrichment independent of chunk composition. The overall distribution of *cat*RAPID ranking scores was right-skewed, with the majority of proteins scoring below 0.55 and a high-confidence tail of interactors exceeding 0.60 **(Figure 2E)**. The 0.55 and 0.60 thresholds shown in **Figure 2A/E** were chosen as descriptive tiers to visually stratify the upper tail of this distribution rather than a statistically derived cutoffs: 0.60 approximates the point beyond which scores fall into the sparsely populated extreme tail (5 of 1,168 proteins, top ∼0.4%), while 0.55 captures a broader “moderate-high” band immediately below it (18 of 1,168 proteins, top ∼1.5%), together restricting the top 1% of the distribution to a score of ≈ 0.58 (12 proteins). These tiers are intended only to help the reader visually distinguish the most extreme scorers in **Figure 2A** from the bulk of the distribution in **Figure 2E**, and carry no independent statistical justification. They are distinct from – and should not be conflated with – the 0.50 threshold used to define the 30 high-confidence interactors analyzed in the sections that follow (**Figures 3–5**); that threshold was derived independently from the hypergeometric enrichment analysis against the experimental RSX interactome (see Methods: Threshold definition for high-confidence interactors). **Supplementary Figure 1** breaks this pattern down by chunk, showing the RNA-binding-domain distribution and top-detected motifs/proteins separately for each of the 9 proteome chunks; the same core interactors (e.g., ELAVL4, AGO1) recur as top hits across multiple chunks despite each chunk containing a distinct protein subset.

**Figure 3.**
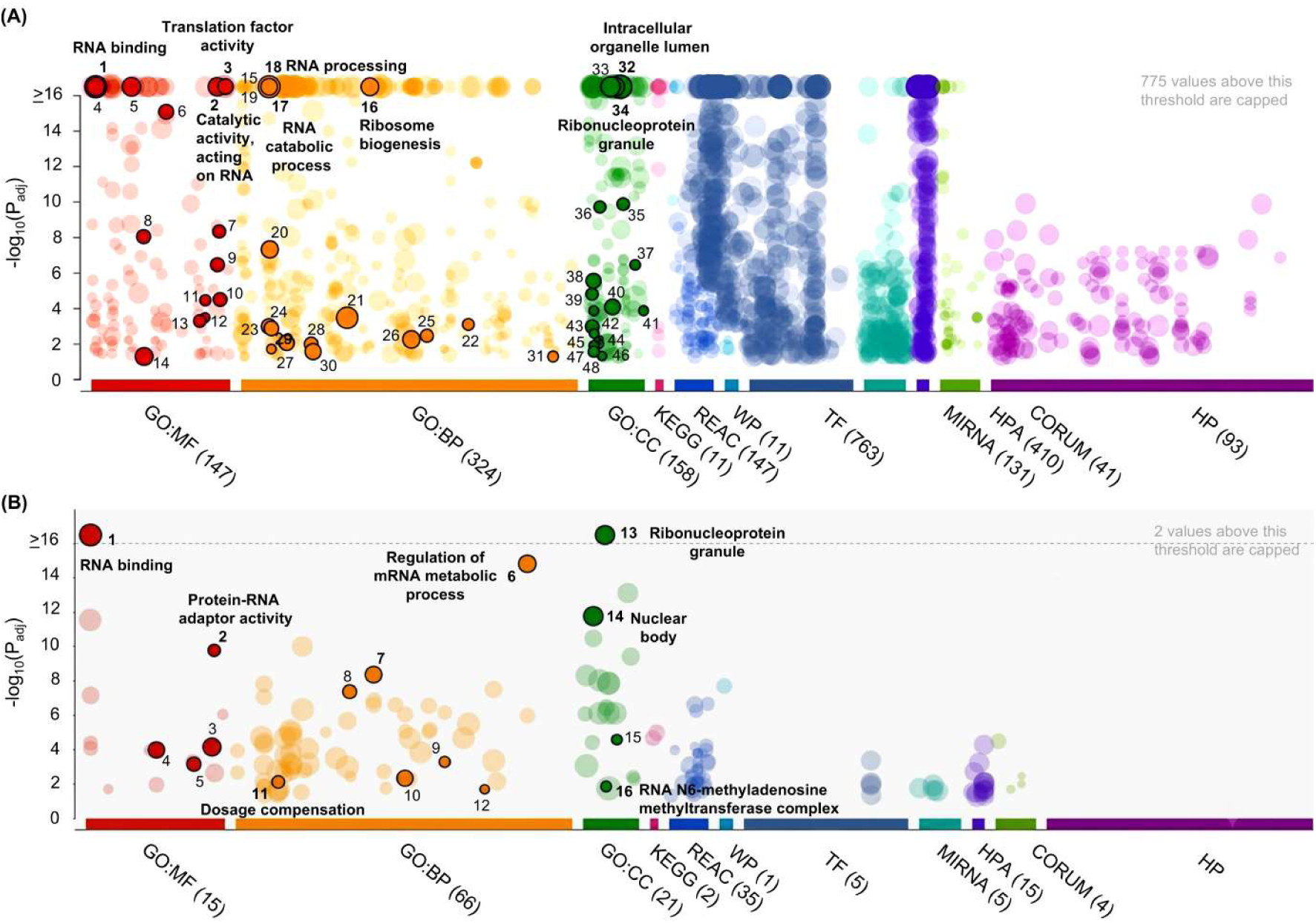
Gene Ontology enrichment analysis of the RSX predicted interactome. Manhattan plots generated by g:Profiler (version e114_eg62_p19; organism: Homo sapiens via human ortholog mapping). The y-axis shows −log₁₀(adjusted p-value); x-axis groups terms by database source. Bubble size reflects gene set size. Highlighted numbered bubbles indicate the most significant non-redundant terms. The dashed line indicates the −log₁₀(p) = 16 cap threshold. Complete results are provided in Supplementary Tables 2–3. **(A)** GO enrichment of the full RSX predicted interactome (1,168 proteins; 2,236 enriched terms; 775 values capped). Top enriched terms include RNA binding (GO:0003723; p = 4.94×10⁻³²⁴), ribonucleoprotein granule (GO:0035770; p = 4.63×10⁻⁹⁸), and RNA processing (p = 3.82×10⁻⁸²), confirming broad enrichment for RNA regulatory and condensate-associated functions. **(B)** GO enrichment of the 30 high-confidence RSX interactors (catRAPID ranking score ≥ 0.50; 169 enriched terms; 2 values capped). Top enriched terms include ribonucleoprotein granule (p = 4.26×10⁻²²), nuclear body (p = 1.69×10⁻¹²), regulation of RNA splicing (p = 4.29×10⁻⁹), and dosage compensation by inactivation of X chromosome (GO:0009048; p = 7.38×10⁻³), the latter driven by METTL3, YTHDC1, and SUZ12.

**Figure 4.**
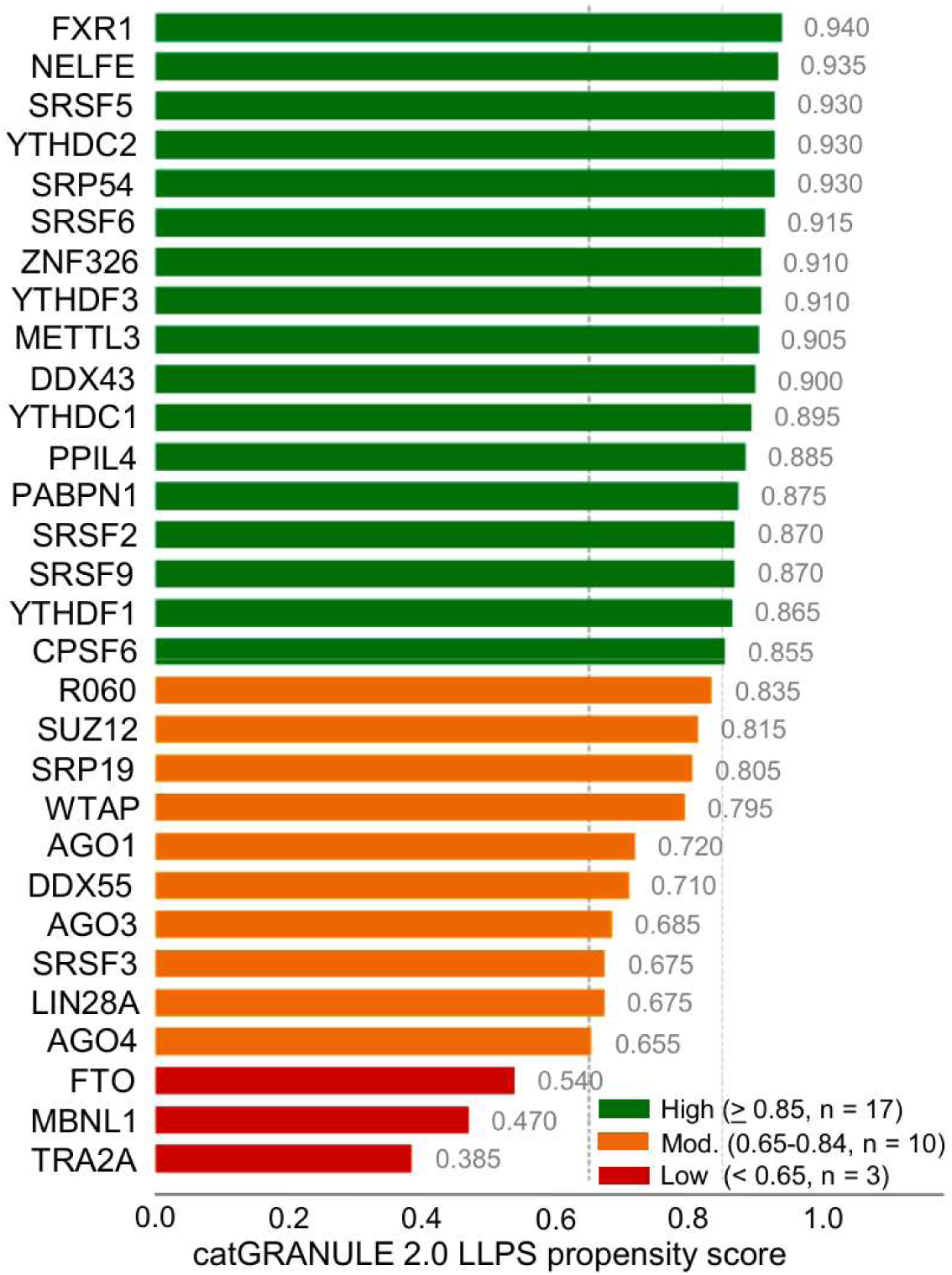
catGRANULE 2.0 liquid-liquid phase separation propensity scores for the 30 high-confidence RSX interactors. Horizontal bar chart showing catGRANULE 2.0 LLPS propensity scores for all 30 high-confidence RSX interactors (catRAPID ranking score ≥ 0.50), ranked from highest (top) to lowest (bottom) score. Scores were computed using native *Monodelphis domestica* protein sequences retrieved from UniProt. Bars are colour-coded by LLPS confidence tier: high LLPS propensity (score ≥ 0.85, green, n = 17), moderate LLPS propensity (0.65–0.84, amber/orange, n = 10), and low LLPS propensity (< 0.65, red, n = 3). The vertical dashed lines indicate the thresholds for moderate (≥ 0.65) and high (≥ 0.85) LLPS confidence. The mean catGRANULE 2.0 score across all 30 proteins (0.803) is indicated by a vertical reference line. Score values are annotated at the end of each bar. The highest-scoring proteins are FXR1 (0.940), NELFE (0.935), YTHDC2 (0.930), SRSF5 (0.930), and SRP54 (0.930). Raw catGRANULE 2.0 scores with UniProt accession numbers are provided in Supplementary Table 4.

**Figure 5.**
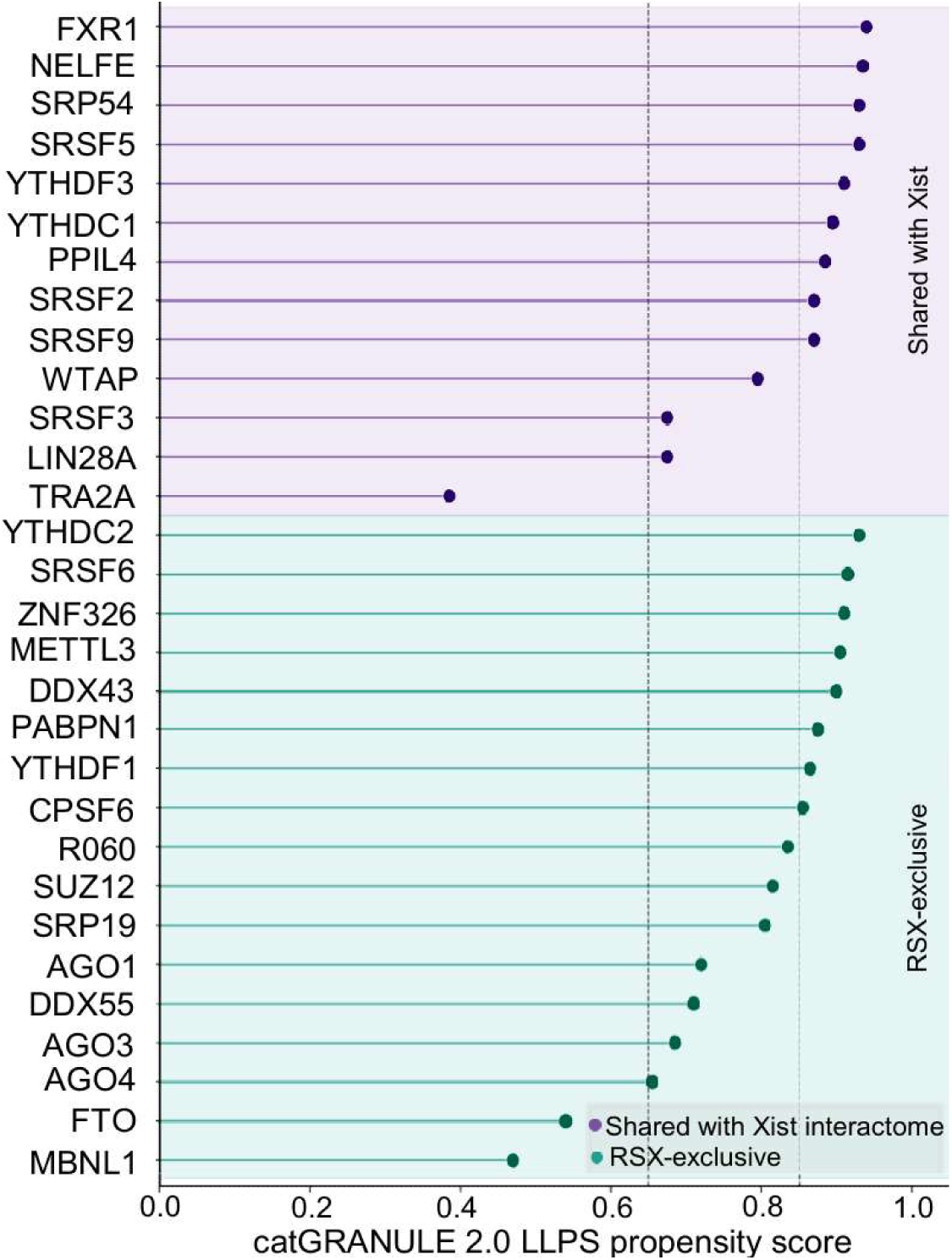
Comparison of the 30 high-confidence RSX interactors with the Xist interactome reveals convergent recruitment of LLPS-competent RNA-binding proteins. Dot/lollipop plot showing catGRANULE 2.0 LLPS propensity scores for all 30 high-confidence RSX interactors, stratified by their relationship to the Xist interactome. Proteins are divided into two categories: shared with the Xist interactome (upper panel, purple, n = 13) and RSX-exclusive (lower panel, teal, n = 17). Within each category, proteins are ranked by LLPS score. Proteins shared with Xist include YTHDC1, YTHDF3, SRSF2, SRSF3, SRSF5, and SRSF9, reflecting convergent recruitment of conserved LLPS-competent RNA-binding machinery. RSX-exclusive proteins with high LLPS scores include ZNF326 (0.910), METTL3 (0.905), DDX43 (0.900), PABPN1 (0.875), and CPSF6 (0.855), suggesting marsupial-specific condensate adaptations.

### Gene Ontology enrichment of the RSX predicted interactome confirms RNA regulatory and condensate-associated functions

To assess the biological coherence of the full predicted RSX interactome, we performed Gene Ontology (GO) enrichment analysis using g:Profiler (version e114_eg62_p19, organism: Homo sapiens via human ortholog mapping) on all 1,168 predicted interactors **(Supplementary Table 2)**. The analysis returned 2,236 significantly enriched terms across Molecular Function (GO:MF), Biological Process (GO:BP), and Cellular Component (GO:CC) categories, reflecting the breadth and functional coherence of the RSX interactome **(Figure 3A)**.

As expected given that the input protein set was drawn from curated RNA-binding protein databases, the most significantly enriched molecular function was RNA binding (GO:0003723; p = 4.94×10⁻³²⁴, 825/1,148 proteins), serving as a technical sanity check on the prediction pipeline rather than a novel biological finding. Catalytic activity acting on RNA (p = 8.60×10⁻²⁰⁰) and translation factor activity (p = 1.05×10⁻⁴⁸) were also prominently enriched, consistent with the diverse post-transcriptional regulatory functions of the predicted interactome. Among biological process terms, translation (p = 4.80×10⁻²³⁸), ribosome biogenesis (p = 1.03×10⁻¹³³), RNA catabolic process (p = 5.02×10⁻⁹⁹), and RNA processing (p = 3.82×10⁻⁸²) were the top enriched categories **(Figure 3A; Supplementary Table 2)**. Critically, cellular component analysis revealed strong enrichment for ribonucleoprotein granule (GO:0035770; p = 4.63×10⁻⁹⁸, 216 proteins) and intracellular organelle lumen (p = 8.79×10⁻¹⁴⁷), providing genome-wide support for the association between RSX interactors and biomolecular condensate compartments **(Figure 3A)**.

GO enrichment of the 30 high-confidence interactors (catRAPID ranking score ≥ 0.50) yielded 169 significantly enriched terms and revealed a highly focused functional signature directly relevant to RSX biology **(Figure 3B; Supplementary Table 3)**. The top cellular component term was ribonucleoprotein granule (p = 4.26×10⁻²², 70 % of the overall proteins), followed by nuclear body (p = 1.69×10⁻¹², 53 %), together providing strong computational evidence that RSX high-confidence interactors are components of phase-separated nuclear condensates **(Figure 3B)**. Among molecular functions, RNA binding (p = 2.94×10⁻¹⁹) and protein-RNA adaptor activity (p = 1.66×10⁻¹⁰) were most enriched. Biological process enrichment highlighted regulation of mRNA metabolic process (p = 1.53×10⁻¹⁵), regulation of RNA splicing (p = 4.29×10⁻⁹), consistent with the recent findings on SR protein-mediated silencing assemblies [34] (**Figure 3**; **Supplementary Table 3**). Strikingly, the GO term "dosage compensation by inactivation of X chromosome" (GO:0009048; p = 7.38×10⁻³) was significantly enriched, driven by METTL3, YTHDC1, and SUZ12 — providing direct ontological support that the predicted RSX interactome captures proteins functionally relevant to X-chromosome inactivation **(Figure 3).** Additionally, the RNA N6-methyladenosine methyltransferase complex (p = 1.33×10⁻²) was enriched, further corroborating the m⁶A regulatory module identified among high-confidence interactors.

### catGRANULE 2.0 LLPS propensity scoring of high-confidence RSX interactors

To assess phase-separation potential with state-of-the-art accuracy, we submitted the 30 high-confidence RSX interactors — using their *Monodelphis domestica* protein sequences retrieved from UniProt — to *cat*GRANULE 2.0, the substantially improved version of the *cat*GRANULE LLPS prediction tool [27] **(Supplementary Table 4)**. Results are summarized in **Table 1** and **Figure 4**.

**Table 1.** Top 30 high-confidence RSX interactors ranked by mean catRAPID ranking with catGRANULE 2.0 LLPS propensity scores. Proteins are ranked by mean catRAPID ranking score aggregated at the gene level across all 9 proteome chunks, with a threshold of catRAPID ranking score ≥ 0.50 defining high-confidence interactors. Columns show: Rank (by mean_ranking); Gene (human ortholog gene symbol used for catRAPID prediction against native *Monodelphis domestica* sequences); Mean Ranking (mean catRAPID ranking score, range 0–1); Max Z-score (maximum interaction Z-score across all proteome chunks); catGRANULE 2.0 LLPS Score (phase separation propensity computed using native M. domestica protein sequences from UniProt); LLPS Confidence tier (High ≥ 0.85, Moderate 0.65–0.84, Low < 0.65). Of 30 proteins, 17 (57%) exhibit high LLPS propensity, 10 moderate, and 3 low. The 3 low-scoring proteins (MBNL1, FTO, TRA2A) represent structurally constrained proteins with limited intrinsic disorder. A broader set of 7 proteins (SRP54, SRP19, SUZ12, TRA2A, MBNL1, FTO, PPIL4) are additionally flagged in Table 1 as non-canonical RBPs, independent of their LLPS score tier.

Of the 30 high-confidence interactors, 17 (57%) exhibited high LLPS propensity (*cat*GRANULE 2.0 score ≥ 0.85; see Methods), 10 exhibited moderate propensity (0.65–0.84), and only 3 scored low (< 0.65) **(Figure 4**; **Table 1)**. The mean *cat*GRANULE 2.0 score across all 30 proteins was 0.803, substantially above background expectation for a random protein set.

The highest-scoring proteins were FXR1 (0.940), NELFE (0.935), YTHDC2 (0.930), SRSF5 (0.930), and SRP54 (0.930) **(Table 1)**. SRP54, a GTPase subunit of the signal recognition particle, harbors a methionine-rich intrinsically disordered M-domain previously implicated in condensate formation [36], consistent with its high catGRANULE 2.0 score of 0.930.

Using a conservative threshold of *ca*tGRANULE 2.0 score ≥ 0.65 (compared to the classifier’s native 0.5 decision boundary) [31], 3 of the 30 high-confidence interactors — TRA2A (0.385), MBNL1 (0.470), and FTO (0.540) — were classified as LLPS-negative. These are either non-canonical RBPs or proteins with well-defined structured domains that limit intrinsic disorder (**Table 1**). These 3 proteins are excluded from the core LLPS narrative while still being noted as predicted interactors.

### Comparison with published Xist interactome datasets: High-confidence RSX interactors are enriched for the m⁶A machinery and splicing regulators

To classify the 30 high-confidence RSX interactors as "shared with Xist" or "RSX-exclusive," each protein was cross-referenced against four independently published experimental Xist interactome datasets: Chu et al. [8], Minajigi et al. [9], McHugh et al. [10], and Cirillo et al. [5,37] (human Xist interactome, included for cross-species comparison). A protein was classified as experimentally shared with Xist if it was reported as a Xist interactor in at least one of the four datasets; proteins absent from all four were classified as RSX-exclusive. The full protein-by-dataset comparison matrix is provided in **Table 2**.

**Table 2.** High-confidence RSX–protein interactors predicted by catRAPID omics v2.0 and their cross-validation against experimentally reported Xist interactomes. Each of the 30 high-confidence RSX interactors (defined by catRAPID ranking score ≥ 0.50; Table 1) is cross-referenced by *Mus musculus* UniProt ID against four independently published Xist interactome datasets: Minajigi et al., Chu et al., McHugh et al., and Cirillo et al. (human). Y indicates the protein was reported as an Xist interactor in that dataset; N indicates it was not. Proteins with at least one "Y" (n = 13) are classified as experimentally shared with the Xist interactome; proteins with "N" across all four datasets (n = 17) are classified as RSX-exclusive/novel candidates.

Among the 17 high-LLPS interactors, two functionally coherent clusters stand out **(Figure 4**; **Table 1)**. First, the m⁶A epitranscriptomic machinery is comprehensively represented: METTL3 (the catalytic methyltransferase, LLPS: 0.905), WTAP (the regulatory subunit, LLPS: 0.795), YTHDC1 (nuclear m⁶A reader, LLPS: 0.895), YTHDC2 (m⁶A helicase, LLPS: 0.930), YTHDF1 (cytoplasmic reader, LLPS: 0.865), and YTHDF3 (cytoplasmic reader, LLPS: 0.910). The co-recruitment of the nuclear writer/reader components (METTL3, WTAP, YTHDC1, YTHDC2) suggests that RSX may nucleate an m⁶A-regulatory condensate on the inactive X; the predicted association with the cytoplasmic readers YTHDF1/YTHDF3 is less readily reconciled with a nuclear silencing model and is addressed further in the discussion.

Second, SR protein splicing regulators form a prominent cluster: SRSF2 (0.870), SRSF3 (0.675), SRSF5 (0.930), SRSF6 (0.915), and SRSF9 (0.870) **(Figure 4**; **Table 1)**. SR proteins are canonical LLPS drivers known to phase-separate into nuclear speckles and regulate co-transcriptional splicing. Their enrichment among RSX interactors parallels observations in the Xist interactome and suggests a conserved role for splicing-factor condensates in XCI.

Additional high-confidence interactors with strong LLPS scores include NELFE (0.935), a negative elongation factor implicated in transcriptional pausing; FXR1 (0.940), an FMR1 homolog involved in translational regulation and stress granule formation; ZNF326 (0.910) and YTHDF3 (0.910), both LLPS-competent nuclear regulators; DDX43 (0.900) and PABPN1 (0.875), helicases and poly(A)-binding proteins with well-established condensate roles **(Table 1)**.

### Comparison with the Xist interactome reveals convergent recruitment of LLPS-competent RBPs

Cross-referencing the 30 high-confidence RSX interactors against four independently published Xist interactome datasets [5,8–10,37] reveals a subset of shared proteins, including members of the YTHDC/YTHDF family, SRSF proteins, and AGO family members (**Figure 5**, **Table 2**). Of the 30 high-confidence interactors, 13 were reported in at least one of the four Xist datasets, while 17 were not detected in any and are classified as RSX-exclusive [8,9]. Proteins interacting exclusively with RSX include ZNF326, METTL3, DDX43, PABPN1 and CPSF6, suggesting marsupial-specific adaptations **(Figure 5**, **Table 2)**. The overlap of functional modules — particularly m⁶A readers and SR splicing factors — despite independent evolutionary origins of RSX and Xist supports convergent recruitment of a conserved LLPS-competent RBP toolkit **(Figure 5)**.

## RSX-Xist conservation and functional coherence

To directly address whether functional convergence between RSX and Xist extends to structural homology, we performed sliding-window minimum free energy (MFE) profiling, contact map analysis, and structural superposition across both transcripts **(Supplementary Figure 2)**. We first computed C3’ atom distance matrices for representative 200-nucleotide windows of each transcript independently, revealing distinct overall fold topologies for RSX and Xist (**Supplementary Figure 2A, B**). Structural superposition of corresponding windows yielded a per-residue RMSD averaging 126.30 Å, with the lowest local divergence at residue 21, indicating a predicted absence of conserved three-dimensional folds between the two lncRNAs (**Supplementary Figure 2C**). Contact maps computed across 200-nucleotide sliding windows further showed largely non-overlapping intramolecular base-pairing patterns for RSX and Xist (**Supplementary Figure 2D, E**); the difference contact map highlighted specific domains of RSX-enriched versus Xist-enriched base-pairing, underscoring divergent internal architectures rather than a shared conserved core (**Supplementary Figure 2F**). Sliding-window MFE profiles for the two transcripts showed only a weak positional correlation (Pearson r = 0.183, p = 0.01), indicating structurally distinct folding landscapes along the length of each RNA (**Supplementary Figure 2G**). Notably, despite this structural divergence, both transcripts showed comparable overall base-pairing propensity (mean paired-base fraction: RSX 55.7 %, Xist 54.0 %), though the positional distribution of paired bases differed substantially between them (**Supplementary Figure 2H**). Collectively, these results confirm that RSX and Xist do not share conserved 2D or 3D folds. These findings complement the sequence-level divergence reported earlier and support a general principle: functional convergence in lncRNA-mediated XCI operates at the level of protein interaction networks and biophysical condensate properties, rather than RNA sequence or structural conservation.

## Discussion

Our comprehensive computational analysis, grounded in native marsupial proteome predictions and rigorously validated against independent experimental data, indicates that RSX preferentially interacts with LLPS-competent RNA-binding proteins enriched in intrinsically disordered regions, supporting the hypothesis that RSX drives XCI through phase separation (**Figures 1**, **4**).The most striking implication of our findings is that RSX and Xist, despite lacking sequence homology and diverging ∼160–180 million years ago, appear to have convergently evolved phase separation as the mechanism for chromosome-wide gene silencing **(Figure 5)**. Several lines of evidence support this convergence. First, top RSX interactors including FXR1, NELFE, and the SRSF family are orthologous to known Xist-binding proteins and play conserved roles in nuclear condensate assembly **(Table 1)**. Second, the mean catGRANULE 2.0 score of 0.803 for RSX’s high-confidence interactors mirrors the LLPS enrichment observed in Xist interactome studies **(Figure 4)**. Third, the comprehensive recruitment of the m⁶A machinery (METTL3, WTAP, YTHDC1/C2, YTHDF1/3) suggests that both lncRNAs may couple epitranscriptomic regulation with phase-separated chromatin compartmentalization (**Table 1**; **Figure 4**), consistent with reports that m⁶A methylation and its WTAP/METTL3/YTHDC1 machinery are directly required for Xist-mediated silencing [19,37]. The additional predicted recruitment of the cytoplasmic readers YTHDF1/YTHDF3 is discussed below as an important caveat of the computational approach. Fourth, the high-confidence enrichment of SR splicing factors (SRSF2, SRSF5, SRSF6, SRSF9) provides a direct parallel to the mechanism recently established by Trotman, Calabrese and colleagues, who showed that Xist Repeat A nucleates an SR-protein assembly to recruit SPEN, with SRSF1 being necessary and sufficient for SPEN recruitment [34]. RSX may employ an analogous, Repeat-A-like mechanism, in which multivalent SR-protein recruitment nucleates condensate assembly and orchestrates gene repression.

It is worth noting that YTHDF1 and YTHDF3 are predominantly cytoplasmic m⁶A readers, in contrast to the nuclear-localized YTHDC1/YTHDC2. Their appearance among high-confidence RSX interactors should therefore be interpreted with caution: catRAPID predictions reflect sequence-level RNA-protein binding compatibility and do not account for subcellular compartmentalization. Direct interaction *in vivo* would require either transient nuclear localization of these readers, co-transcriptional engagement prior to RSX’s presumed nuclear retention, or reflect residual cytoplasmic RSX transcript pools; this distinction is not resolved by the present computational approach and represents an important caveat for future experimental validation.

Existing imaging data further aligns with this physical model. Conventional RNA-FISH imaging of RSX shows a compact, granular signal within the Xi territory [11], qualitatively similar to the punctate appearance of Xist RNA clouds under standard confocal resolution. While consistent with a biomolecular condensate framework, diffraction-limited imaging cannot resolve true nanoscale condensate architecture; super-resolution approaches (e.g., 3D-SIM, STORM), previously applied to Xist [38], have not yet been applied to RSX and represent an essential next step to provide direct structural validation.

The GO enrichment results provide independent biological validation of both the full interactome and the high-confidence subset **(Figures 3A, B; Supplementary Tables 2–3)**. The enrichment of ribonucleoprotein granule and nuclear body terms among the top 30 interactors (p = 4.26×10⁻²² and p = 1.69×10⁻¹², respectively) independently corroborates the catGRANULE 2.0 LLPS scores and confirms that these proteins are established components of phase-separated nuclear compartments. Most compellingly, the direct enrichment of the GO term "dosage compensation by inactivation of X chromosome" among the top 30 predicted interactors — driven by METTL3, YTHDC1, and SUZ12 **(Figure 3B)** — provides ontological validation that our computational predictions are capturing functionally relevant biology rather than non-specific RNA-binding associations. This convergence of interaction prediction, LLPS scoring, and GO annotation on XCI-relevant biology substantially strengthens confidence in the predicted RSX interactome.

The convergent evolution of LLPS-based XCI in marsupials and eutherians suggests that phase separation provides fundamental biophysical advantages for chromosome-scale regulation that natural selection has independently arrived at twice. That RSX and Xist independently exploit these properties underscores LLPS as an evolutionarily optimal solution to the dosage compensation problem. Notably, similar phase-separation mechanisms have also been implicated in Drosophila dosage compensation and heterochromatin organization, suggesting LLPS may represent a universal principle for chromosome-scale gene regulation across metazoans [39–41].

## Conclusions

Our integrated computational framework demonstrates that marsupial RSX and eutherian Xist—despite ∼160–180 million years of divergence, absence of sequence homology, and distinct 2D/3D topologies—convergently recruit an orthologous, LLPS-competent RBP toolkit enriched in m⁶A machinery and SR splicing factors. These findings indicate that liquid-liquid phase separation and epitranscriptomic compartmentalization represent evolutionarily conserved biophysical strategies for chromosome-scale gene silencing across mammalian lineages. This study establishes a validated RBP network and provides a clear roadmap for experimental testing of RSX condensates in marsupial cell models (**Figure 6**).

**Figure 6.**
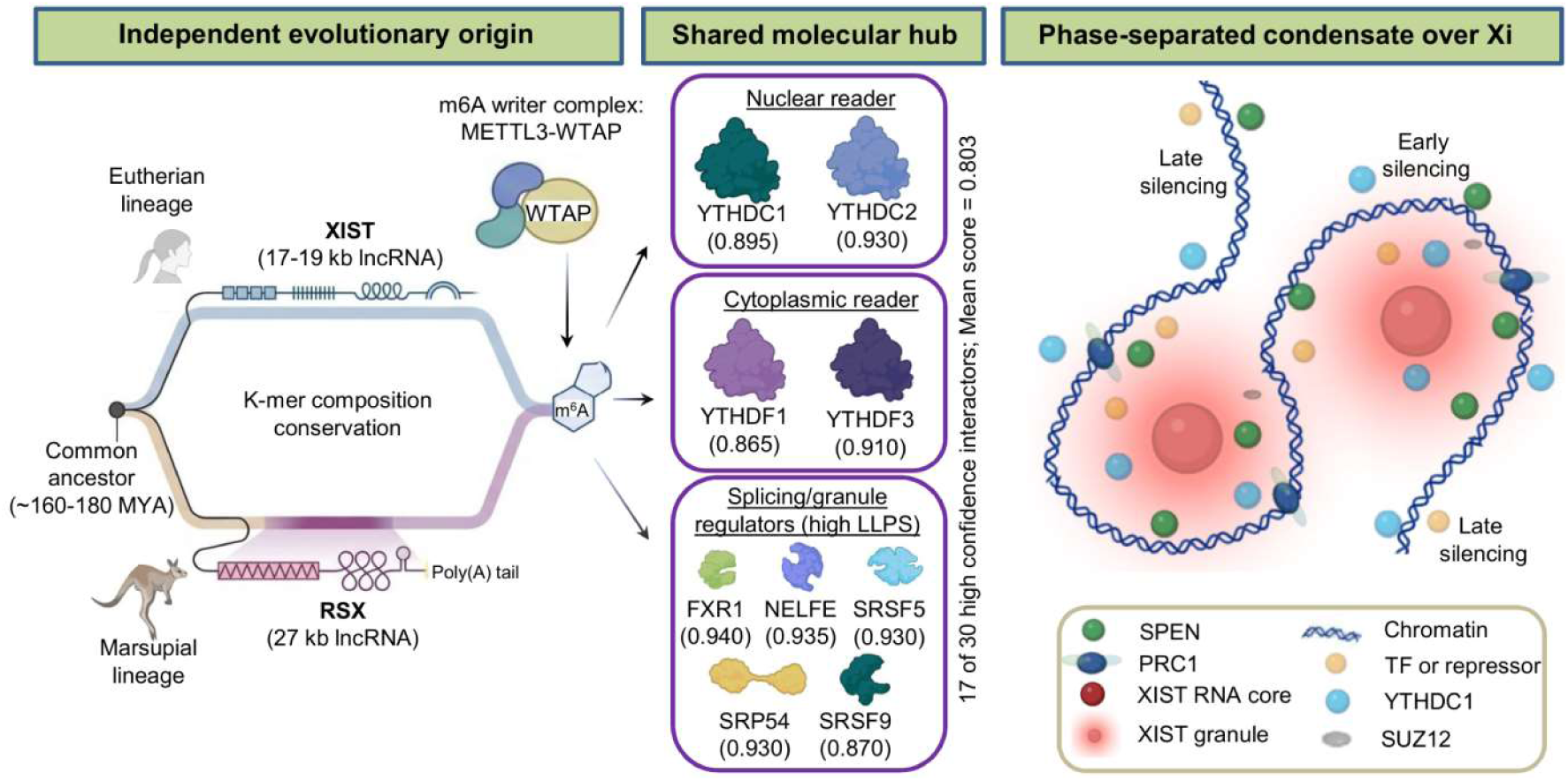
Convergent RSX and Xist biology despite independent evolutionary origins. RSX (27 kb; marsupial lineage) and Xist (17–19 kb; eutherian lineage) diverged from a common ancestor ∼160– 180 million years ago and share no sequence homology or conserved secondary/tertiary fold (per-residue RMSD > 126 Å), despite conserved k-mer composition. Both lncRNAs converge on a shared hub of LLPS-competent RNA-binding proteins centred on the m⁶A machinery the METTL3–WTAP writer complex, nuclear readers YTHDC1/YTHDC2, and cytoplasmic readers YTHDF1/YTHDF3 together with high-LLPS splicing and granule regulators (FXR1, NELFE, SRSF5, SRP54, SRSF9). These represent a subset of the 30 high-confidence RSX interactors identified by catRAPID omics v2.0 (mean = 0.803), of which 17 (57%) show high LLPS propensity (*cat*GRANULE 2.0 score ≥ 0.85; mean = 0.901). In eutherians, Xist nucleates a phase-separated condensate over the inactive X (Xi), excluding transcriptional machinery and recruiting established silencing factors (SPEN/SHARP, Polycomb) alongside METTL3, SUZ12, and YTHDC1. Whether RSX forms an analogous condensate over Xi is a model proposed here, pending experimental confirmation.

## Methods

### Statistical validation against the experimental RSX interactome

To assess the predictive performance of catRAPID, predictions were validated against an independently published experimental RSX interactome [33]. All catRAPID prediction files were merged and unique proteins extracted, yielding a prediction universe of 1,140 proteins. Experimentally identified RSX interactors from the reference dataset were matched to the prediction universe (47 matched proteins; baseline probability 4.1%). Proteins were ranked by mean catRAPID ranking at the gene level, and enrichment of experimental hits among top-ranked predictions was assessed using hypergeometric tests. Enrichment analyses were performed at top 50 and top 100 thresholds, and separately for proteins shared between RSX and Xist experimental datasets.

### Expected-by-chance calculation

The experimental Rsx interactome (13059_2024_3280_MOESM2_ESM.csv) was matched against the 1,140-protein prediction universe, yielding 47 matched proteins and a baseline recovery probability of 47/1,140 ≈ 0.041 (4.1%) for a random draw. The number of experimental hits expected by chance among the top N predictions was calculated as this baseline probability multiplied by N (e.g., 0.041 × 50 ≈ 2.06 for the top 50; 0.041 × 100 ≈ 4.12 for the top 100). Enrichment was then calculated as observed/expected hits, and statistical significance assessed by the hypergeometric test (comparing observed overlap against the full 1,140-protein universe, 47 successes, N draws).

### Threshold definition for high-confidence interactors

A threshold of catRAPID ranking score ≥ 0.50 was applied to define high-confidence RSX interactors. This threshold was selected based on the statistical enrichment analysis: it corresponds to the point of strongest enrichment of experimentally validated proteins and is supported by the hypergeometric significance of the enrichment at multiple thresholds (top 50: p ≈ 1.6×10⁻⁵; top 100: p ≈ 9.5×10⁻⁵). At this threshold, 30 proteins were identified, of which 13 are experimentally supported and 17 are novel candidates.

### catRAPID omics v2.0 predictions against the *Monodelphis domestica* proteome

The full-length RSX RNA sequence (27 kb) from *Monodelphis domestica* (accession JQ937282.1) was analyzed using the *catRAPID omics* v2.0 server (http://service.tartaglialab.com/page/catrapid_omics2_group) [28,30]. To ensure biological consistency, predictions were performed against the *Monodelphis domestica* RBP proteome rather than the human proteome. The complete proteome was submitted in 9 validated chunks to ensure full coverage, yielding 1,168 unique proteins. catRAPID predicts RNA-protein interaction propensities based on physicochemical parameters including secondary structure, hydrogen bonding, and van der Waals forces. Each RNA-protein pair is assigned a Z-score and a ranking score. Proteins were aggregated at the gene level using mean ranking across all RNA fragments and proteome chunks.

### Gene Ontology enrichment analysis

GO term enrichment was performed using g:Profiler (version e114_eg62_p19; https://biit.cs.ut.ee/gprofiler/gost) [42] with Homo sapiens as the reference organism via human ortholog mapping, due to limited *Monodelphis domestica* annotation in g:Profiler databases. Enrichment significance was assessed using g:SCS multiple testing correction. Analysis was performed separately for the full 1,168-protein predicted interactome and the 30 high-confidence interactors. Enriched terms across GO Biological Process (GO:BP), Molecular Function (GO:MF), Cellular Component (GO:CC), KEGG, Reactome, and additional annotation databases were reported. Terms with adjusted p < 0.05 were considered significant.

### catGRANULE 2.0 LLPS propensity scoring

For the 30 high-confidence RSX interactors (catRAPID ranking score ≥ 0.50), LLPS propensity scores were computed using *cat*GRANULE 2.0 (https://tools.tartaglialab.com/catgranule2) [31]. Protein sequences were retrieved from UniProt for *Monodelphis domestica* (taxon ID: 13616); where multiple isoforms existed, the longest canonical isoform was used. catGRANULE 2.0 computes LLPS potential using improved sequence-derived features including intrinsically disordered regions (IDRs), low-complexity domains, prion-like motifs, and RNA-binding domains. Scores range from 0 (low) to 1 (high); proteins with scores ≥ 0.85 were classified as high LLPS propensity, 0.65–0.84 as moderate, and < 0.65 as low. The 0.65 threshold reflects a conservative offset from catGRANULE 2.0’s native 0.5 decision boundary. The 0.85 cutoff was applied only to visually stratify the highest-scoring candidates within this dataset and does not reflect a natural discontinuity in the score distribution or an independently validated boundary; all enrichment statistics and conclusions are based on continuous scores rather than the categorical tiers.

### RNA secondary structure analysis

Secondary structure analysis of RSX (*Monodelphis domestica*; accession JQ937282) and human Xist (accession NR_001564) was performed using the ViennaRNA package (v2.6.x) [43]. Minimum free energy (MFE) secondary structures were predicted using RNAfold in sliding windows of 200 nucleotides with a step size of 100 nucleotides across the full length of each transcript. Pseudo-three-dimensional coordinates were generated from dot-bracket secondary structures, and pairwise distances were computed using the C3’ ribose backbone atom as the structural reference point, analogous to the Cα atom convention used in protein structural comparison. Per-window MFE values and paired-base fractions were extracted and compared between transcripts using Pearson correlation. Contact maps were constructed from dot-bracket structure outputs by recording all base-paired positions within each window and aggregating across the transcript. Difference contact maps were computed by subtracting Xist contact frequencies from RSX contact frequencies at equivalent relative positions. Structural superposition of representative RSX and Xist window structures, based on C3’ atom coordinates, was performed using the US-align algorithm [44], and per-residue RMSD values were extracted from superposition outputs.

### Statistical analyses

Hypergeometric tests were used to assess enrichment of experimentally validated proteins among top-ranked catRAPID predictions. All statistical analyses were performed in Python (v3.12) using SciPy. A significance threshold of p < 0.05 was applied.

### Data visualization

catRAPID omics v2.0 interaction plots were generated directly by the catRAPID server. LLPS score distribution and RSX versus Xist comparison plots were generated using Matplotlib and Seaborn in Python 3.12. GO enrichment Manhattan plots were generated by g:Profiler. Data processing was performed using Pandas, NumPy, and Matplotlib libraries

## Abbreviations

XCI: X-chromosome inactivation
LLPS: Liquid-liquid phase separation
lncRNA: Long non-coding RNA
RSX: RNA on the silent X
Xist: X-inactive specific transcript
Xi: Inactive X chromosome
RBP: RNA-binding protein
IDR: Intrinsically disordered region
GO: Gene Ontology
FISH: Fluorescence in situ hybridization
RIP: RNA immunoprecipitation
MFE: Minimum free energy

## Declarations

### Ethics approval and consent to participate

Not applicable (computational study only).

### Consent for publication

Not applicable.

### Availability of data and materials

All data supporting the conclusions of this article are included within the article and its supplementary files. The RSX RNA sequence used for predictions is publicly available under GenBank accession JQ937282.1. The *Monodelphis domestica* proteome used for catRAPID omics v2.0 predictions is publicly available from UniProt (taxon ID: 13616). catGRANULE 2.0 LLPS propensity scores were computed using the catGRANULE 2.0 server (https://tools.tartaglialab.com/catgranule2). Gene Ontology enrichment analyses were performed using g:Profiler (https://biit.cs.ut.ee/gprofiler/gost; version e114_eg62_p19).

The catRAPID omics v2.0 prediction results for all 9 *Monodelphis domestica* proteome chunks are publicly accessible via the following job links:

- Chunk 0 (mondo.RBP.0.fasta): http://service.tartaglialab.com/email_redir/1138545/2db9eb0ac2
- Chunk 1 (mondo.RBP.1.fasta): http://service.tartaglialab.com/email_redir/1138553/98f18de7ba
- Chunk 2 (mondo.RBP.2.fasta): http://service.tartaglialab.com/email_redir/1138554/37d3202361
- Chunk 3 (mondo.RBP.3.fasta): http://service.tartaglialab.com/email_redir/1138555/dde824e8c4
- Chunk 4 (mondo.RBP.4.fasta): http://service.tartaglialab.com/email_redir/1138556/d5ead5b48a
- Chunk 5 (mondo.RBP.5.fasta): http://service.tartaglialab.com/email_redir/1138557/286d02312e
- Chunk 6 (mondo.RBP.6.fasta): http://service.tartaglialab.com/email_redir/1138558/3d000014fc
- Chunk 7 (mondo.RBP.7.fasta): http://service.tartaglialab.com/email_redir/1138559/9db4ca5ee3
- Chunk 8 (mondo.RBP.8.fasta): http://service.tartaglialab.com/email_redir/1138560/6fdad9473a

The complete predicted interactome, top 30 high-confidence interactor scores, catGRANULE 2.0 LLPS scores, and GO enrichment results are provided as Supplementary Tables 1–4. Custom analysis scripts used for statistical validation and data processing are available from the corresponding author upon reasonable request.

Note: catRAPID server job links are provided for transparency; raw prediction data are fully available in Supplementary Table 1 to ensure long-term accessibility.

### Authors’ contributions

AKD and CC contributed equally to this work. AKD, CC, RDP, and GGT performed computational analyses, data visualization, and contributed to manuscript writing. AC conceived and supervised the study, interpreted results, and wrote the manuscript. All authors read and approved the final manuscript.

## Acknowledgements

We thank the developers of catRAPID omics v2.0, catGRANULE 2.0, and g:Profiler for maintaining publicly accessible computational resources that made this analysis possible. We acknowledge Google Colab for providing computational resources used in data analysis and visualization.

## Table legends

**Supplementary Table 1. Complete catRAPID omics v2.0 predicted RSX interactome against the *Monodelphis domestica* proteome.** Full prediction output for all 1,168 unique proteins across 9 proteome chunks. Columns include:Protein_ID (gene name with GN. prefix), RNA_ID, Top_RNA_fragment (nucleotide positions of the highest-scoring RSX fragment), Annotation, Interaction_Propensity, Z_score, RBP_Propensity, RNA_Binding_Domains_IDs (Pfam domain identifiers), Number of RNA-Binding Domain Instances, RNA_Binding_Motifs_IDs, Number of RNA-Binding Motif Instances, and Ranking score. Proteins are listed in descending order of Ranking score. Note: the merged and deduplicated prediction universe used for statistical validation comprised 1,140 unique proteins after cross-chunk integration.

**Supplementary Table 2. Gene Ontology enrichment results for the full RSX predicted interactome (1,168 proteins).** Complete g:Profiler enrichment output (version e114_eg62_p19; organism: Homo sapiens) for all 1,168 predicted RSX interactors. Columns include: source (GO:MF, GO:BP, GO:CC, KEGG, REAC, WP, TF, MIRNA, HPA, CORUM, HP), term_name, term_id, highlighted (Boolean, indicating terms selected for Manhattan plot labelling), adjusted_p_value (g:SCS multiple testing correction), −log₁₀(adjusted p-value), term_size, query_size, intersection_size, effective_domain_size, and intersections (gene symbols driving enrichment). A total of 2,236 significantly enriched terms are reported. Analysis was performed using human ortholog gene symbols due to limited *Monodelphis domestica* annotation in g:Profiler databases.

**Supplementary Table 3. Gene Ontology enrichment results for the 30 high-confidence RSX interactors.** Complete g:Profiler enrichment output (version e114_eg62_p19; organism: Homo sapiens) for the 30 high-confidence RSX interactors (catRAPID ranking score ≥ 0.50). Columns as described in Supplementary Table 2. A total of 169 significantly enriched terms are reported. Key terms include ribonucleoprotein granule (GO:0035770; p = 4.26×10⁻²²), nuclear body (p = 1.69×10⁻¹²), RNA binding (p = 2.94×10⁻¹⁹), regulation of RNA splicing (p = 4.29×10⁻⁹), and dosage compensation by inactivation of X chromosome (GO:0009048; p = 7.38×10⁻³), the latter driven by METTL3, YTHDC1, and SUZ12.

**Supplementary Table 4. catGRANULE 2.0 LLPS propensity scores for 30 high-confidence RSX interactors using native *Monodelphis domestica* protein sequences.** Raw catGRANULE 2.0 output for all 30 high-confidence RSX interactors. Columns include: Rank (by mean catRAPID ranking), Gene (human ortholog symbol), UniProt accession (native *M. domestica* sequence), species confirmation (*Monodelphis_domestica*), and catGRANULE 2.0 LLPS propensity score (range 0–1). All sequences were retrieved directly from UniProt using native *M. domestica* accession numbers to ensure species-appropriate LLPS scoring. Scores ≥ 0.85 indicate high phase separation propensity; 0.65–0.84 moderate; < 0.65 low.

## References

1. Lyon MF. Gene Action in the X-chromosome of the Mouse (Mus musculus L.). Nature. Nature Publishing Group; 1961;190:372–3. 10.1038/190372a0

2. Galupa R, Heard E. X-Chromosome Inactivation: A Crossroads Between Chromosome Architecture and Gene Regulation. Annu Rev Genet. 2018;52:535–66. 10.1146/annurev-genet-120116-024611

3. Brown CJ, Ballabio A, Rupert JL, Lafreniere RG, Grompe M, Tonlorenzi R, et al. A gene from the region of the human X inactivation centre is expressed exclusively from the inactive X chromosome. Nature. Nature Publishing Group; 1991;349:38–44. 10.1038/349038a0

4. Brockdorff N, Ashworth A, Kay GF, McCabe VM, Norris DP, Cooper PJ, et al. The product of the mouse Xist gene is a 15 kb inactive X-specific transcript containing no conserved ORF and located in the nucleus. Cell. 1992;71:515–26. 10.1016/0092-8674(92)90519-I

5. Cerase A, Armaos A, Neumayer C, Avner P, Guttman M, Tartaglia GG. Phase separation drives X-chromosome inactivation: a hypothesis. Nat Struct Mol Biol. 2019;26:331–4. 10.1038/s41594-019-0223-0

6. Pandya-Jones A, Markaki Y, Serizay J, Chitiashvili T, Mancia Leon WR, Damianov A, et al. A protein assembly mediates Xist localization and gene silencing. Nature. 2020;587:145–51. 10.1038/s41586-020-2703-0

7. Perotti I, Broglia L, Tartaglia GG, Cerase A. Xist condensates: perspectives for therapeutic intervention. Genome Biol. 2025;26:215. 10.1186/s13059-025-03666-8

8. Chu C, Zhang QC, da Rocha ST, Flynn RA, Bharadwaj M, Calabrese JM, et al. Systematic Discovery of Xist RNA Binding Proteins. Cell. 2015;161:404–16. 10.1016/j.cell.2015.03.025

9. Minajigi A, Froberg J, Wei C, Sunwoo H, Kesner B, Colognori D, et al. Chromosomes. A comprehensive Xist interactome reveals cohesin repulsion and an RNA-directed chromosome conformation. Science. New York, N.Y.; 2015;349:10.1126/science.aab2276 aab2276. 10.1126/science.aab2276

10. McHugh CA, Chen C-K, Chow A, Surka CF, Tran C, McDonel P, et al. The Xist lncRNA interacts directly with SHARP to silence transcription through HDAC3. Nature. Nature Publishing Group; 2015;521:232–6. 10.1038/nature14443

11. Grant J, Mahadevaiah SK, Khil P, Sangrithi MN, Royo H, Duckworth J, et al. Rsx is a metatherian RNA with Xist-like properties in X-chromosome inactivation. Nature. 2012;487:254–8. 10.1038/nature11171

12. Duret L, Chureau C, Samain S, Weissenbach J, Avner P. The *Xist* RNA Gene Evolved in Eutherians by Pseudogenization of a Protein-Coding Gene. Science. 2006;312:1653–5. 10.1126/science.1126316

13. Sprague D, Waters SA, Kirk JM, Wang JR, Samollow PB, Waters PD, et al. Nonlinear sequence similarity between the Xist and Rsx long noncoding RNAs suggests shared functions of tandem repeat domains. RNA. New York, N.Y.; 2019;25:1004–19. 10.1261/rna.069815.118

14. Whitworth DJ, Pask AJ. The X factor: X chromosome dosage compensation in the evolutionarily divergent monotremes and marsupials. Semin Cell Dev Biol. 2016;56:117–21. 10.1016/j.semcdb.2016.01.006

15. Renfree MB, Hore TA, Shaw G, Marshall Graves JA, Pask AJ. Evolution of Genomic Imprinting: Insights from Marsupials and Monotremes. Annu Rev Genomics Hum Genet. 2009;10:241–62. 10.1146/annurev-genom-082908-150026

16. Chaumeil J, Waters PD, Koina E, Gilbert C, Robinson TJ, Marshall Graves JA. Evolution from XIST-Independent to XIST-Controlled X-Chromosome Inactivation: Epigenetic Modifications in Distantly Related Mammals. Feil R, editor. PLoS ONE. 2011;6:e19040. 10.1371/journal.pone.0019040

17. Mahadevaiah SK, Sangrithi MN, Hirota T, Turner JMA. A single-cell transcriptome atlas of marsupial embryogenesis and X inactivation. Nature. 2020;586:612–7. 10.1038/s41586-020-2629-6

18. Hornecker JL, Samollow PB, Robinson ES, VandeBerg JL, McCarrey JR. Meiotic sex chromosome inactivation in the marsupial *Monodelphis domestica*. genesis. 2007;45:696–708. 10.1002/dvg.20345

19. Patil DP, Chen C-K, Pickering BF, Chow A, Jackson C, Guttman M, et al. m6A RNA methylation promotes XIST-mediated transcriptional repression. Nature. 2016;537:369–73. 10.1038/nature19342

20. Ries RJ, Zaccara S, Klein P, Olarerin-George A, Namkoong S, Pickering BF, et al. m6A enhances the phase separation potential of mRNA. Nature. 2019;571:424–8. 10.1038/s41586-019-1374-1

21. Banani SF, Lee HO, Hyman AA, Rosen MK. Biomolecular condensates: organizers of cellular biochemistry. Nat Rev Mol Cell Biol. 2017;18:285–98. 10.1038/nrm.2017.7

22. Shin Y, Brangwynne CP. Liquid phase condensation in cell physiology and disease. Science. 2017;357:eaaf4382. 10.1126/science.aaf4382

23. Guo YE, Manteiga JC, Henninger JE, Sabari BR, Dall’Agnese A, Hannett NM, et al. Pol II phosphorylation regulates a switch between transcriptional and splicing condensates. Nature. 2019;572:543–8. 10.1038/s41586-019-1464-0

24. Yamazaki T, Souquere S, Chujo T, Kobelke S, Chong YS, Fox AH, et al. Functional Domains of NEAT1 Architectural lncRNA Induce Paraspeckle Assembly through Phase Separation. Mol Cell. 2018;70:1038–1053.e7. 10.1016/j.molcel.2018.05.019

25. Colognori D, Sunwoo H, Kriz AJ, Wang C-Y, Lee JT. Xist Deletional Analysis Reveals an Interdependency between Xist RNA and Polycomb Complexes for Spreading along the Inactive X. Mol Cell. 2019;74:101–117.e10. 10.1016/j.molcel.2019.01.015

26. Pintacuda G, Wei G, Roustan C, Kirmizitas BA, Solcan N, Cerase A, et al. hnRNPK Recruits PCGF3/5-PRC1 to the Xist RNA B-Repeat to Establish Polycomb-Mediated Chromosomal Silencing. Mol Cell. Elsevier; 2017;68:955–969.e10. 10.1016/j.molcel.2017.11.013

27. Monfort A, Di Minin G, Postlmayr A, Freimann R, Arieti F, Thore S, et al. Identification of Spen as a Crucial Factor for Xist Function through Forward Genetic Screening in Haploid Embryonic Stem Cells. Cell Rep. 2015;12:554–61. 10.1016/j.celrep.2015.06.067

28. Armaos A, Colantoni A, Proietti G, Rupert J, Tartaglia GG. catRAPID omics v2.0: going deeper and wider in the prediction of protein-RNA interactions. Nucleic Acids Res. 2021;49:W72–9. 10.1093/nar/gkab393

29. Bolognesi B, Faure AJ, Seuma M, Schmiedel JM, Tartaglia GG, Lehner B. The mutational landscape of a prion-like domain. Nat Commun. 2019;10:4162. 10.1038/s41467-019-12101-z

30. Bellucci M, Agostini F, Masin M, Tartaglia GG. Predicting protein associations with long noncoding RNAs. Nat Methods. 2011;8:444–5. 10.1038/nmeth.1611

31. Monti M, Fiorentino J, Miltiadis-Vrachnos D, Bini G, Cotrufo T, Sanchez de Groot N, et al. catGRANULE 2.0: accurate predictions of liquid-liquid phase separating proteins at single amino acid resolution. Genome Biol. 2025;26:33. 10.1186/s13059-025-03497-7

32. Bolognesi B, Lorenzo Gotor N, Dhar R, Cirillo D, Baldrighi M, Tartaglia GG, et al. A Concentration-Dependent Liquid Phase Separation Can Cause Toxicity upon Increased Protein Expression. Cell Rep. 2016;16:222–31. 10.1016/j.celrep.2016.05.076

33. McIntyre KL, Waters SA, Zhong L, Hart-Smith G, Raftery M, Chew ZA, et al. Identification of the RSX interactome in a marsupial shows functional coherence with the Xist interactome during X inactivation. Genome Biol. 2024;25:134. 10.1186/s13059-024-03280-0

34. Trotman JB, Porrello A, Schactler SA, DeLeon LE, Eberhard QE, Boyson SP, et al. Xist Repeat A coordinates an assembly of SR proteins to recruit SPEN and induce gene silencing. bioRxiv: The Preprint Server for Biology; 2025. 10.1101/2025.05.21.655143

35. Cirillo D, Blanco M, Armaos A, Buness A, Avner P, Guttman M, et al. Quantitative predictions of protein interactions with long noncoding RNAs. Nat Methods. 2016;14:5–6. 10.1038/nmeth.4100

36. Mateju D, Franzmann TM, Patel A, Kopach A, Boczek EE, Maharana S, et al. An aberrant phase transition of stress granules triggered by misfolded protein and prevented by chaperone function. EMBO J. 2017;36:1669–87. 10.15252/embj.201695957

37. Moindrot B, Cerase A, Coker H, Masui O, Grijzenhout A, Pintacuda G, et al. A Pooled shRNA Screen Identifies Rbm15, Spen, and Wtap as Factors Required for Xist RNA-Mediated Silencing. Cell Rep. 2015;12:562–72. 10.1016/j.celrep.2015.06.053

38. Smeets D, Markaki Y, Schmid VJ, Kraus F, Tattermusch A, Cerase A, et al. Three-dimensional super-resolution microscopy of the inactive X chromosome territory reveals a collapse of its active nuclear compartment harboring distinct Xist RNA foci. Epigenetics Chromatin. 2014;7:8. 10.1186/1756-8935-7-8

39. Strom AR, Emelyanov AV, Mir M, Fyodorov DV, Darzacq X, Karpen GH. Phase separation drives heterochromatin domain formation. Nature. 2017;547:241–5. 10.1038/nature22989

40. Larson AG, Elnatan D, Keenen MM, Trnka MJ, Johnston JB, Burlingame AL, et al. Liquid droplet formation by HP1α suggests a role for phase separation in heterochromatin. Nature. 2017;547:236–40. 10.1038/nature22822

41. Erdel F, Rippe K. Formation of Chromatin Subcompartments by Phase Separation. Biophys J. 2018;114:2262–70. 10.1016/j.bpj.2018.03.011

42. Raudvere U, Kolberg L, Kuzmin I, Arak T, Adler P, Peterson H, et al. g:Profiler: a web server for functional enrichment analysis and conversions of gene lists (2019 update). Nucleic Acids Res. 2019;47:W191–8. 10.1093/nar/gkz369

43. Lorenz R, Bernhart SH, Höner Zu Siederdissen C, Tafer H, Flamm C, Stadler PF, et al. ViennaRNA Package 2.0. Algorithms Mol Biol AMB. 2011;6:26. 10.1186/1748-7188-6-26

44. Zhang C, Shine M, Pyle AM, Zhang Y. US-align: universal structure alignments of proteins, nucleic acids, and macromolecular complexes. Nat Methods. 2022;19:1109–15. 10.1038/s41592-022-01585-1

