## Supplementary figures and images for "Phase Separation Potential of Marsupial RSX RNA Reveals Convergent Evolution of X-Chromosome Inactivation Mechanisms"

### Supplemetary Figure 1

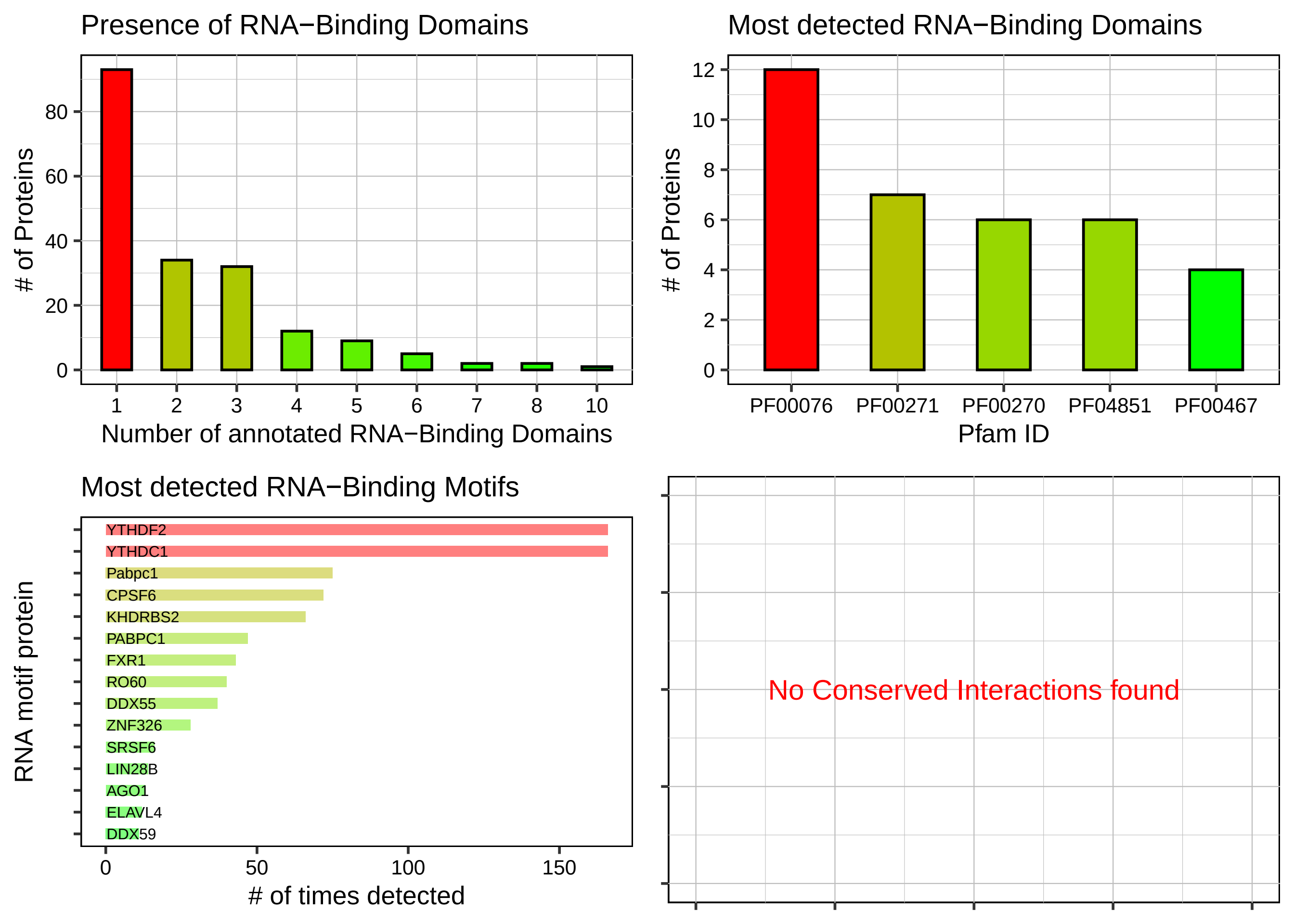

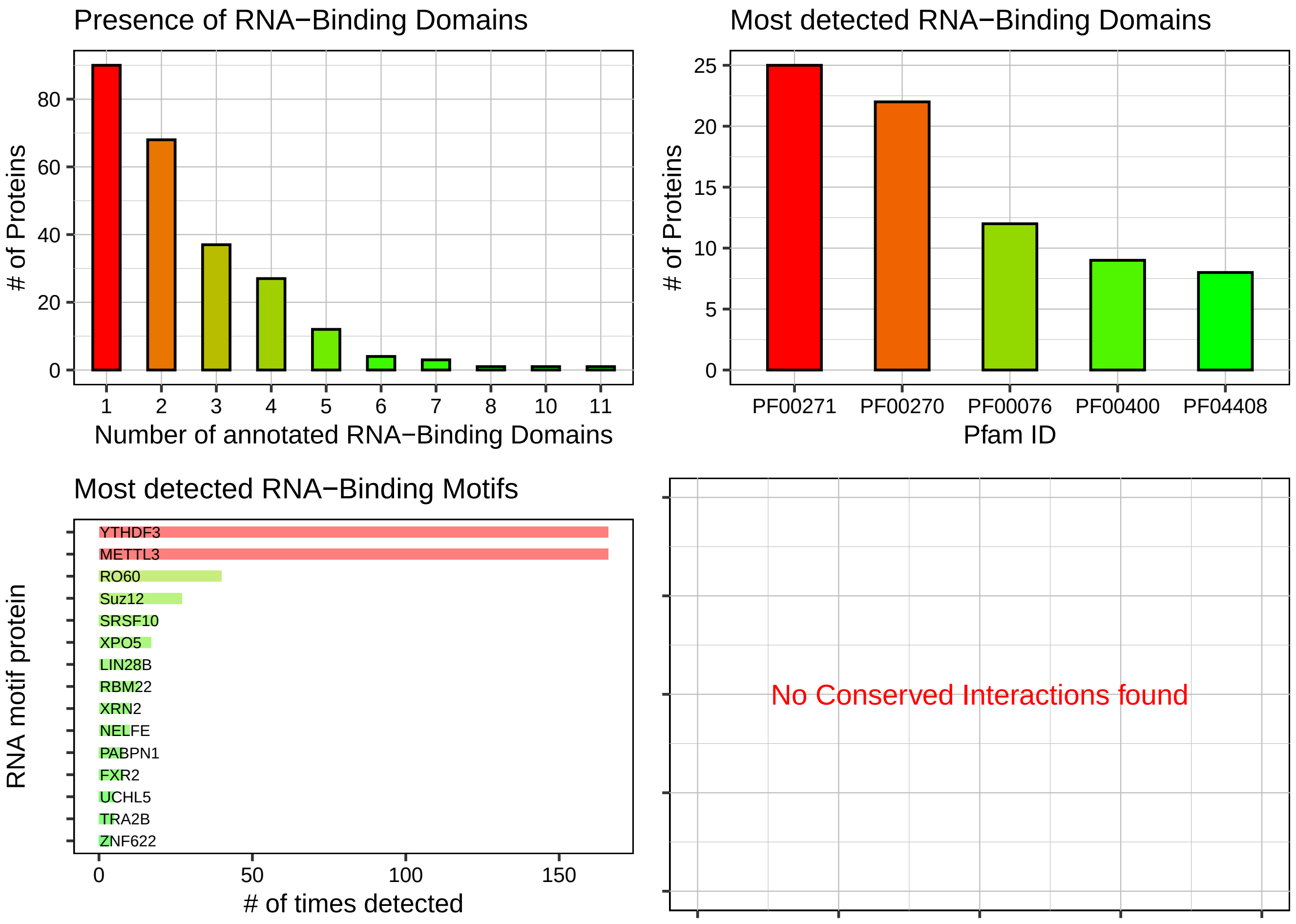

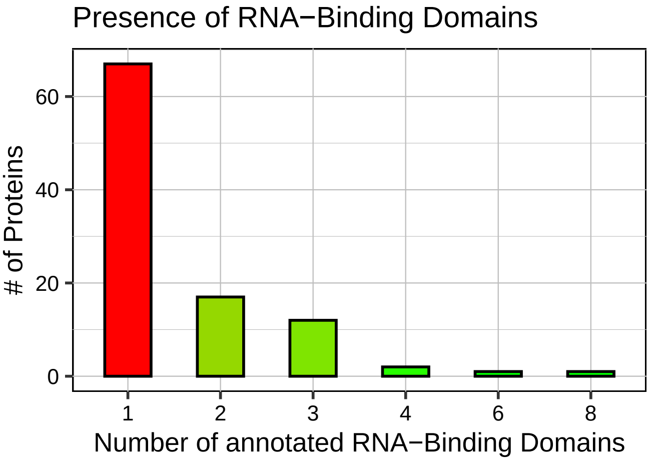

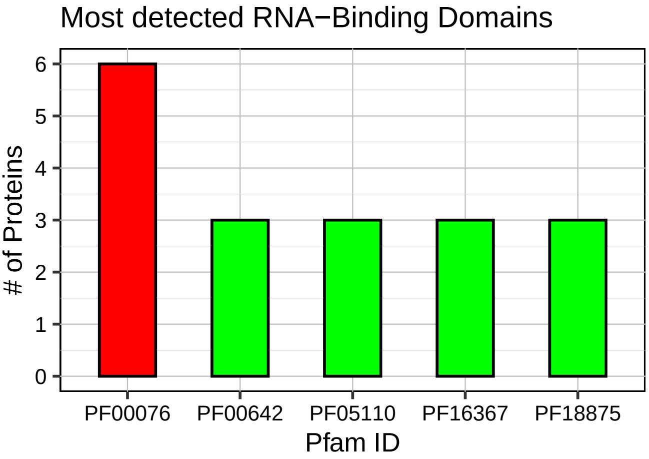

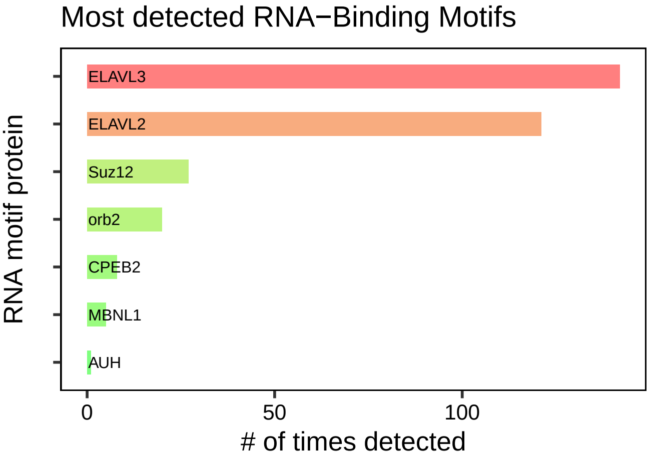

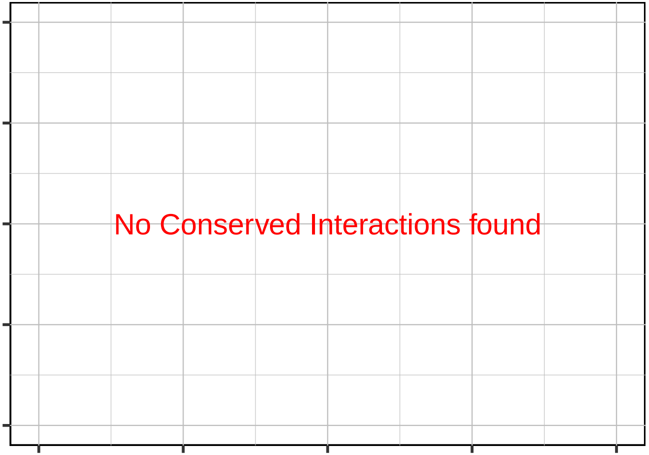

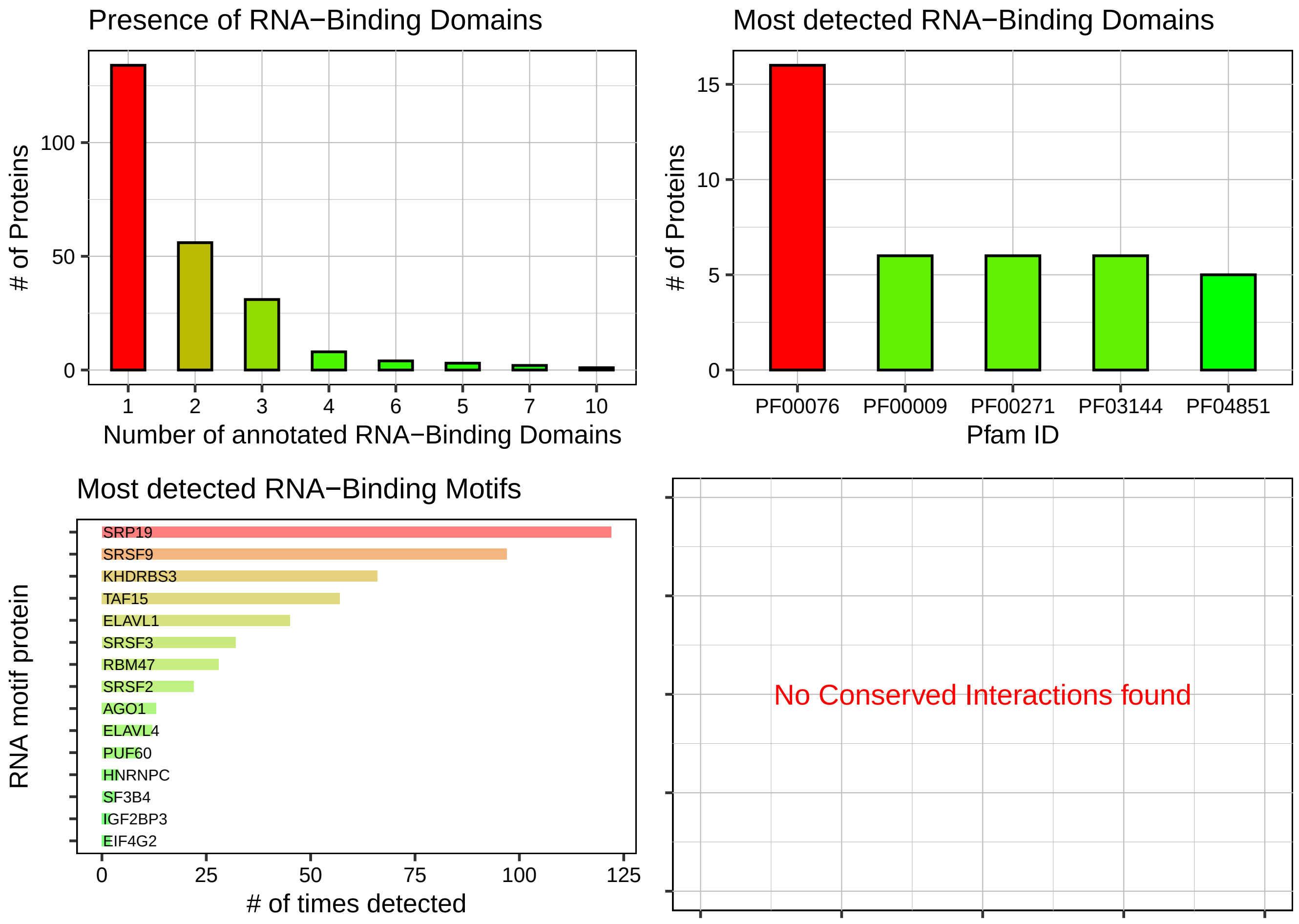

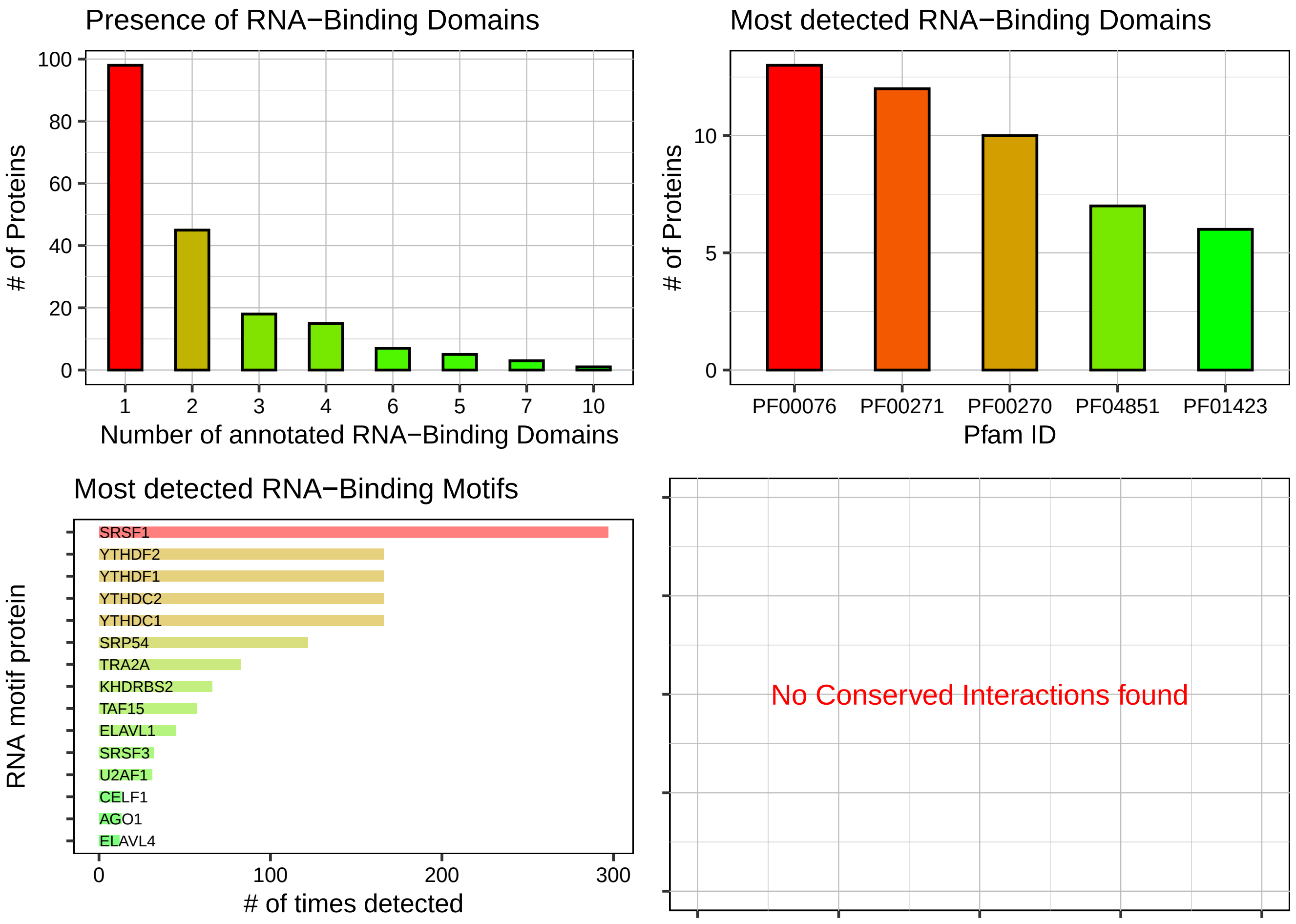

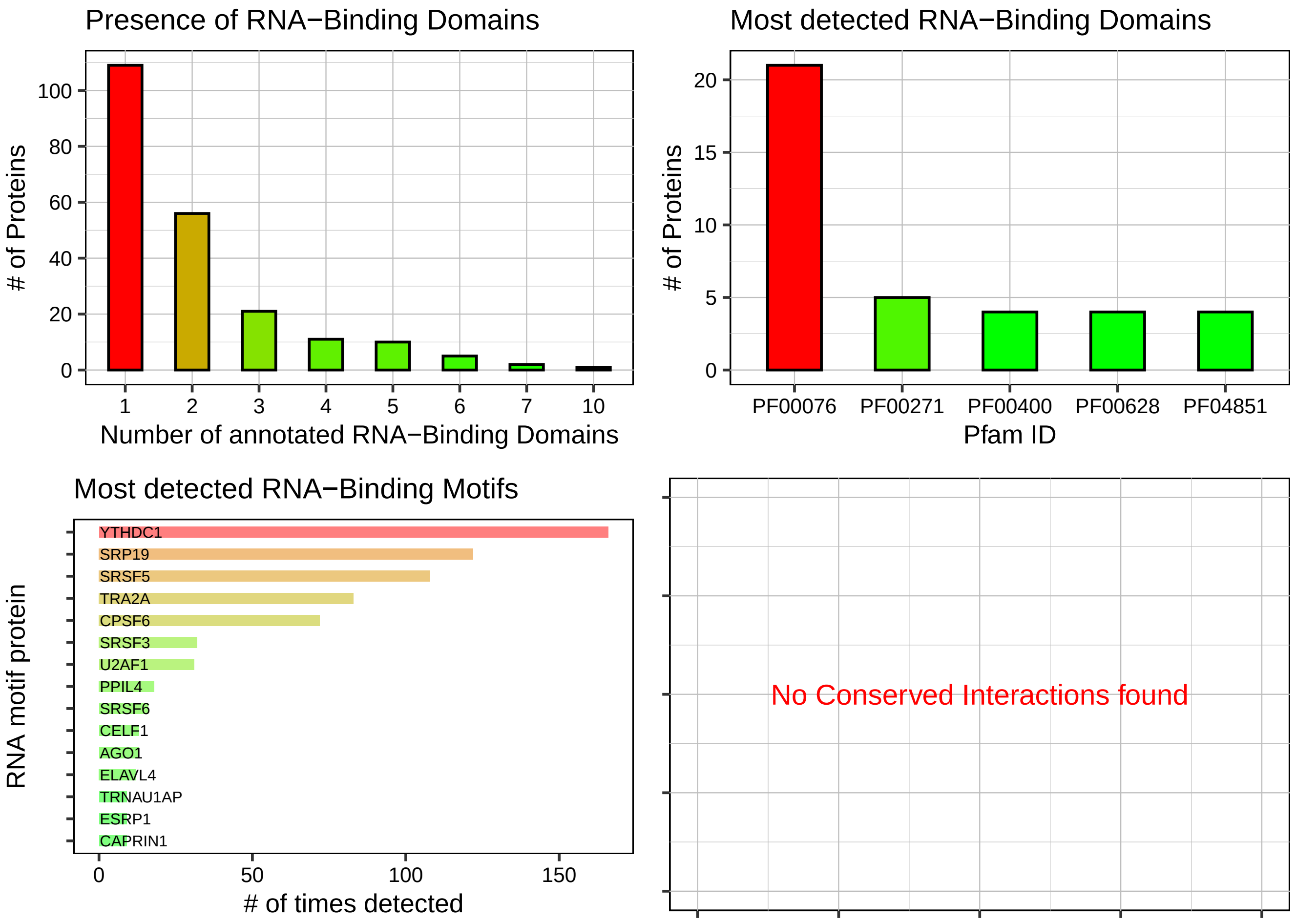

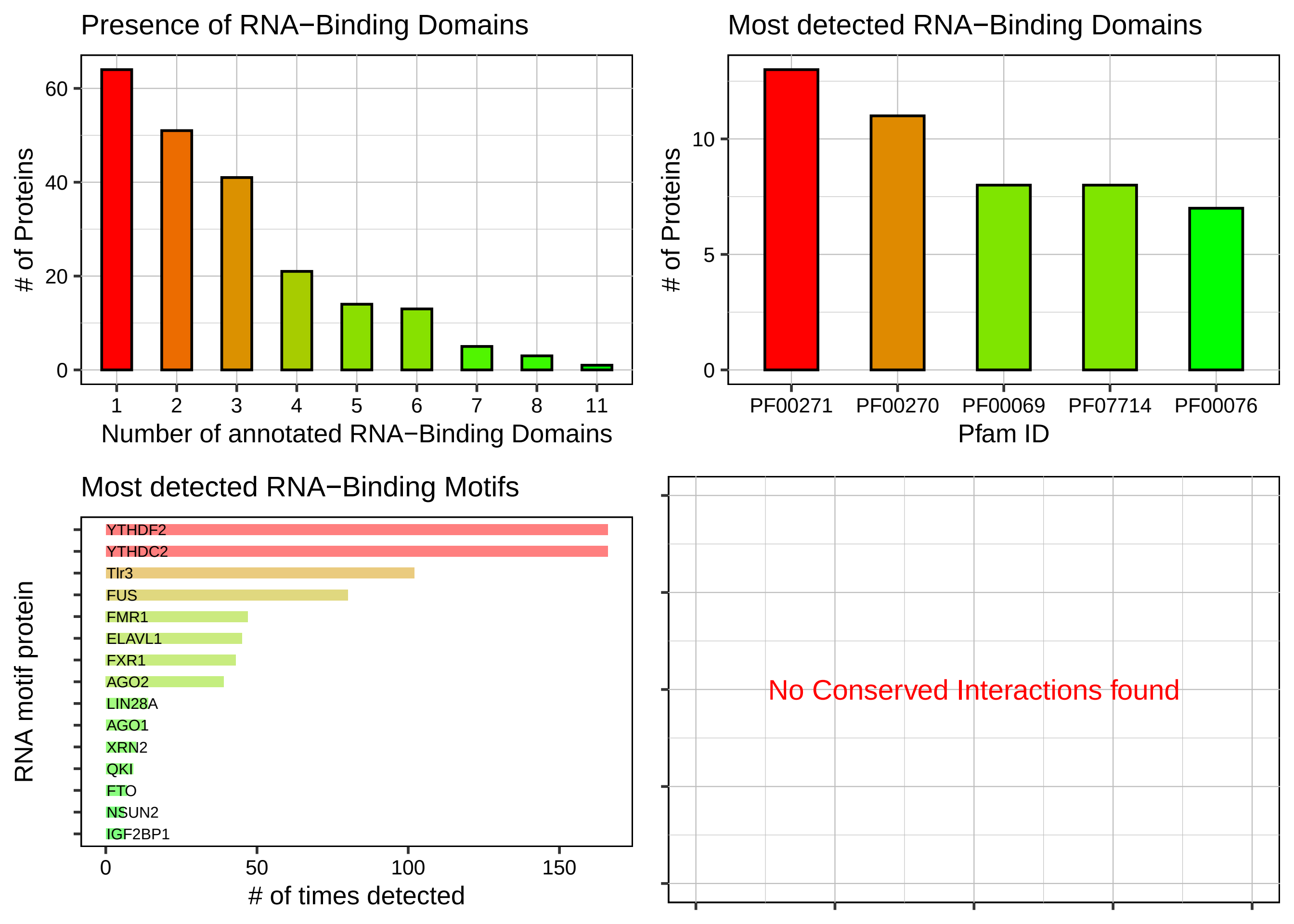

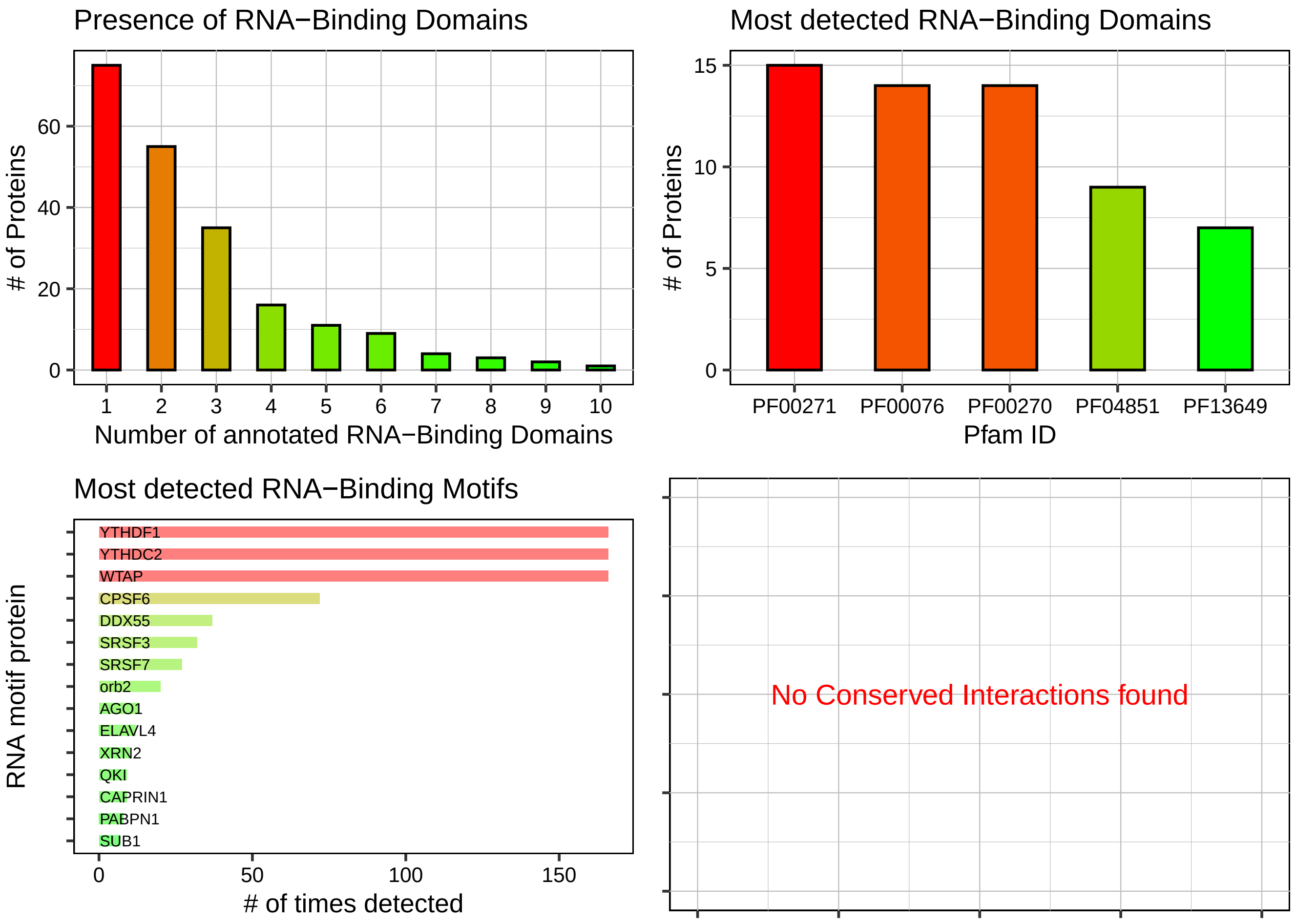

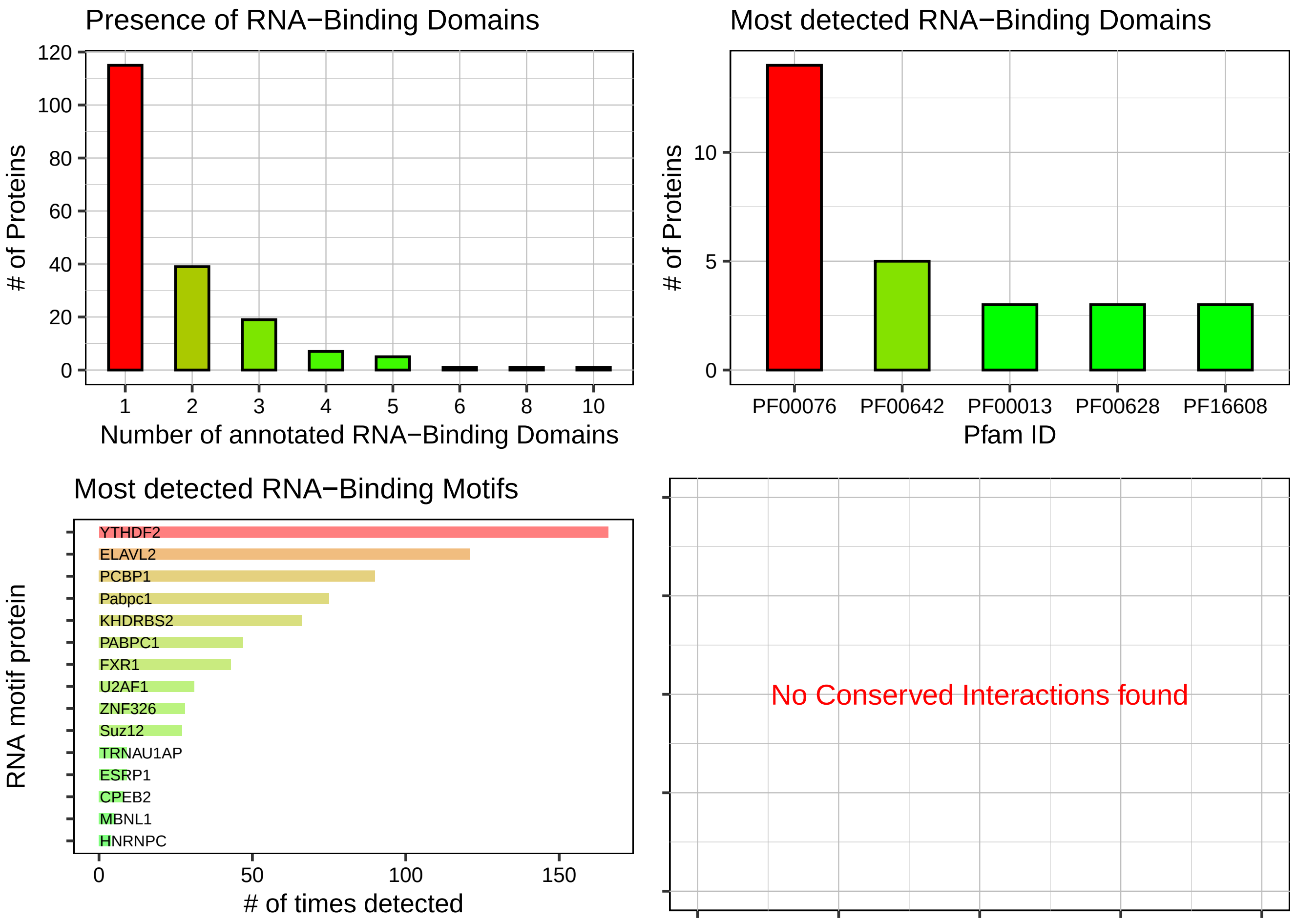
