## Supplementary material for "Phase Separation Potential of Marsupial RSX RNA Reveals Convergent Evolution of X-Chromosome Inactivation Mechanisms": Supplemetary Figure 2

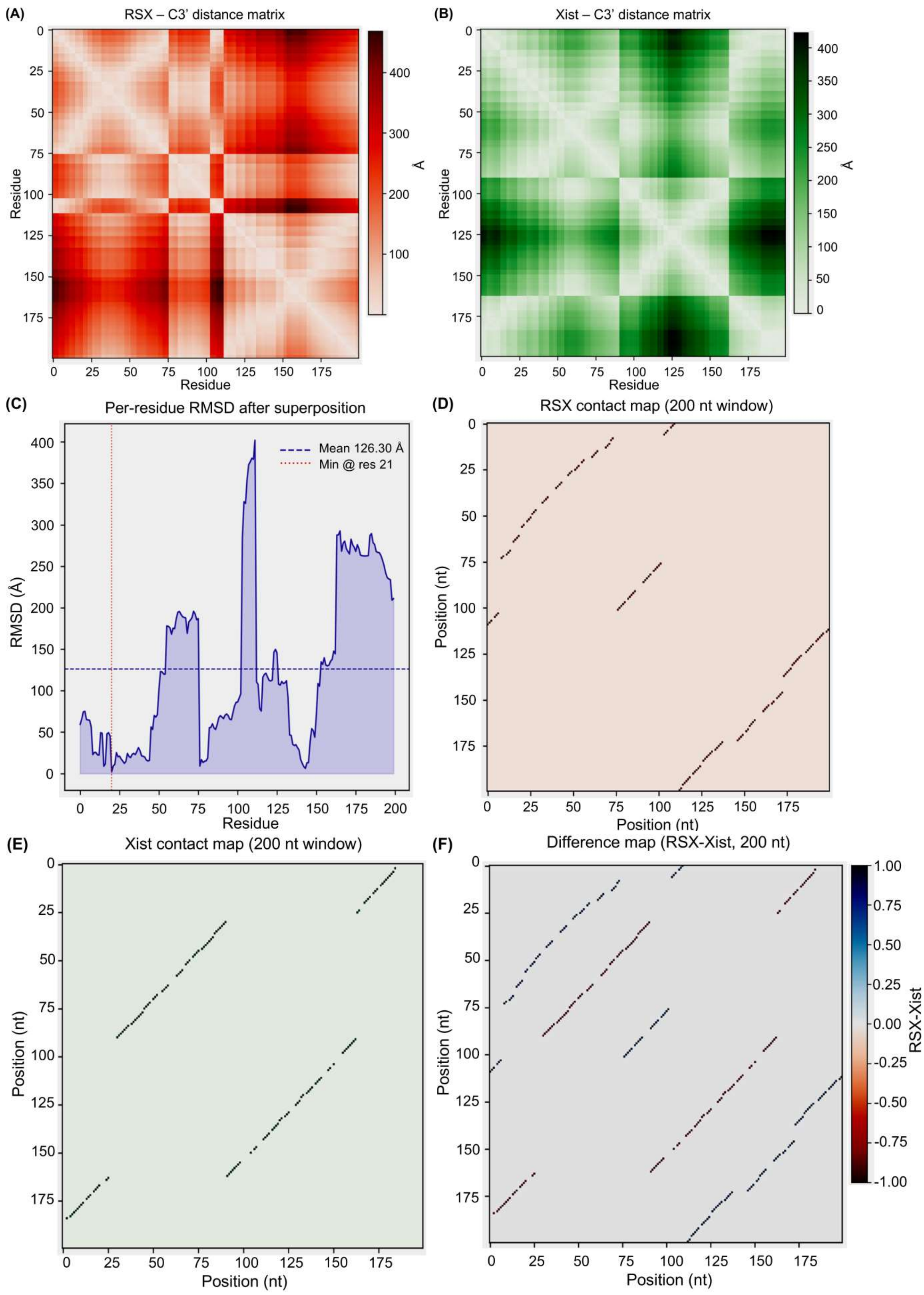
(A)
RSX – C3' distance matrix
Residue
Å
(B)
Xist – C3' distance matrix
Residue
Å
(C)
Per-residue RMSD after superposition
Mean 126.30 Å
Min @ res 21
RMSD (Å)
Residue
(D)
RSX contact map (200 nt window)
Position (nt)
(E)
Xist contact map (200 nt window)
Position (nt)
(F)
Difference map (RSX-Xist, 200 nt)
RSX-Xist
Position (nt)

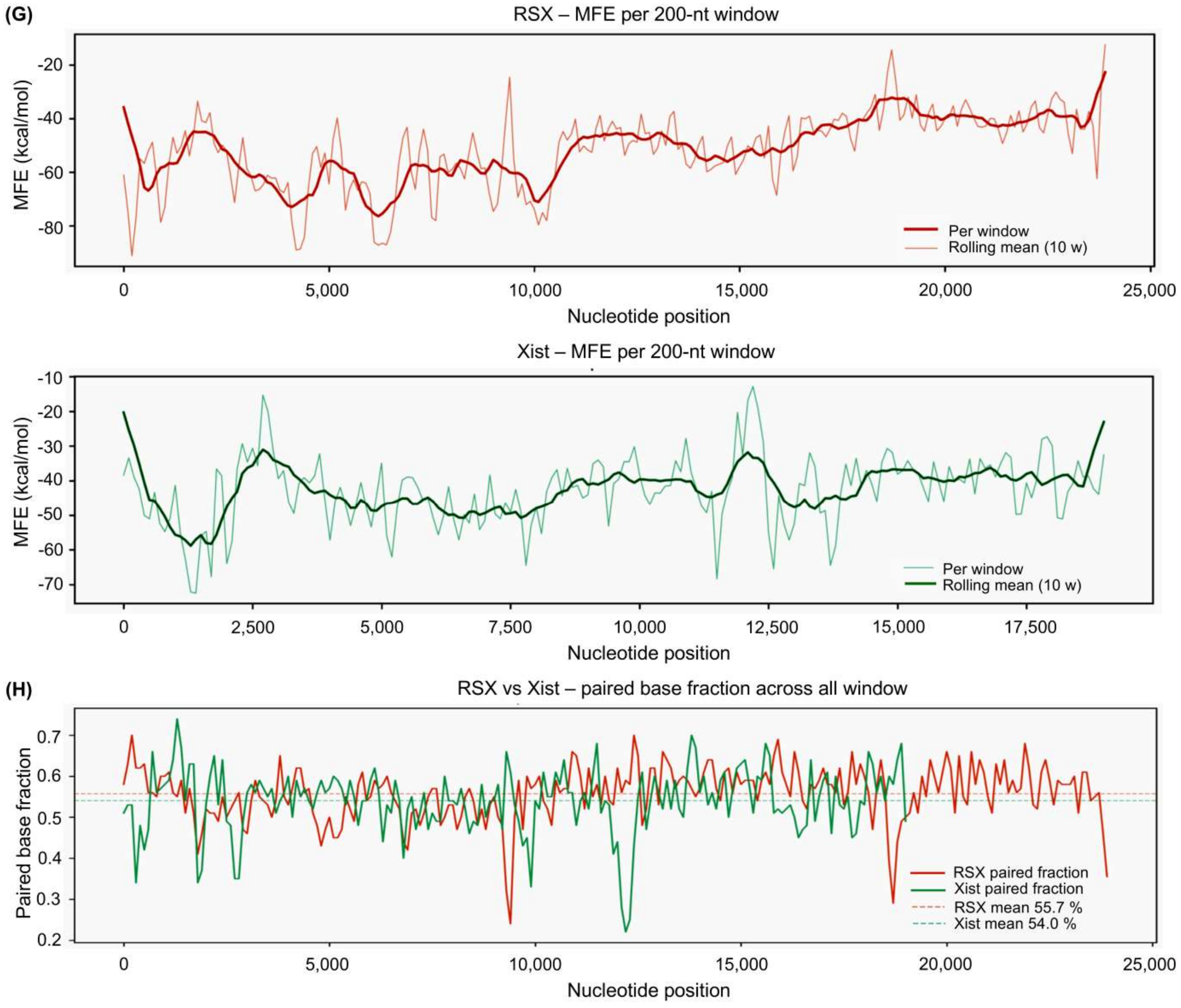


**Supplementary Figure 2. RSX and Xist lack conserved secondary structure despite similar base-pairing propensity. (A, B)** C3' backbone-atom distance matrices computed for representative 200-nucleotide windows of RSX (A, red) and Xist (B, green), illustrating the overall fold topology of each transcript independently. **(C)** Per-residue RMSD following structural superposition of RSX and Xist windows (mean RMSD = 126.30 Å; minimum at residue 21), confirming the absence of conserved three-dimensional folds between the two transcripts. **(D, E)** Contact maps computed for 200-nucleotide sliding windows of RSX (D) and Xist (E), representing intramolecular base-pairing interactions across each transcript. **(F)** Difference contact map (RSX minus Xist) highlighting divergent domain architectures between the two lncRNAs, with positive values (red) indicating RSX-enriched contacts and negative values (blue) indicating Xist-enriched contacts. **(G)** Sliding-window minimum free energy (MFE) profiles for RSX (red) and Xist (green) computed in 200-nucleotide windows across the full length of each transcript, with 10-window rolling means shown as solid lines. Pearson correlation between positional MFE profiles was weak (r = 0.183, p = 0.01), indicating structurally distinct folding landscapes. **(H)** Paired-base fraction profiles across all sliding windows for RSX (red, mean = 55.7%) and Xist (green, mean = 54.0%), shown with mean values indicated by dashed vertical lines. Despite comparable overall base-pairing propensity, the positional distribution of paired bases differed substantially between transcripts.
